# Teaching Diffusion Models Physics: Reinforcement Learning for Physically Valid Diffusion-Based Docking

**DOI:** 10.64898/2026.03.25.714128

**Authors:** J. Henry Broster, Bojana Popovic, Diana Kondinskaia, Charlotte M. Deane, Fergus Imrie

## Abstract

Molecular docking aims to predict the binding conformation of a small molecule to its protein target. Recent work has proposed diffusion models for this task, from rigid-body docking that diffuses over ligand degrees of freedom to co-folding approaches that jointly generate protein structure and ligand pose. However, diffusion-based docking models have been shown to frequently produce physically implausible poses and fail to consistently recover key protein-ligand interactions. To address this, we introduce a reinforcement learning framework for training diffusion-based docking models directly on non-differentiable objectives. Fine-tuning DiffDock-Pocket for physical validity with our approach substantially increases the number of generated poses that are physically valid and interaction-preserving, with no increase in inference-time compute. Importantly, this comes without sacrificing structural accuracy; in fact, our approach increases the proportion of structures with near-native poses. These effects are most pronounced for protein targets that are dissimilar to the training data. Our fine-tuned DiffDock-Pocket model outperforms both classical docking algorithms and machine learning-based approaches on the PoseBusters set. Our results demonstrate that reinforcement learning can teach diffusion-based docking models to better respect physical constraints and recover key interactions, without the requirement to rely on inference-time corrections.

## 1 Introduction

Drug discovery remains slow, costly, and marked by high late-stage attrition [1, 2]. This motivates computational methods that can more reliably prioritize candidates before expensive experimental and clinical testing [1]. Structure-based drug design (SBDD) exploits three-dimensional (3D) information about the target and ligand structures to identify, design, and optimize candidate molecules [3]. A key component of SBDD is molecular docking, which predicts the ligand orientation and conformation in the binding site and is widely used for virtual screening and lead optimization [4, 5]. Docking can reduce experimental burden by prioritizing compounds more likely to succeed, but in practice prospective hit rates are often low and can vary substantially across targets and screening setups [6–9].

Classical approaches to molecular docking rely on physics-inspired scoring functions combined with heuristic or stochastic search algorithms. Docking tools such as AutoDock Vina [10], Glide [11], and GOLD [12] generate poses by searching over the translational, torsional, and rotational space that the ligand can take using, for instance, a genetic algorithm before ranking them using physics-based scoring functions, which serve as a proxy for binding free energy. More recently, machine learning models have increasingly been adopted to replace either the search or scoring components of docking. Early machine learning (ML)-based approaches, such as GNINA [13], used 3D convolutional neural networks to rescore poses generated with classical algorithms, while models such as EquiBind [14] and TankBind [15] operate directly in 3D space to predict ligand poses in a single forward pass. These models are trained with supervised regression objectives, producing a single pose prediction that may average over multiple valid binding modes rather than capturing any one accurately. As a result, they do not model the inherent multimodality of protein-ligand binding, motivating the use of diffusion-based generative models, which have been shown to better model multimodal distributions [16–18].

Diffusion generative models [19, 20] are a class of machine learning models that learn complex data distributions by reversing a stochastic noise process. Diffusion models define a forward process that gradually corrupts data with noise and learn a reverse process that iteratively removes this noise, generating new samples from the learned distribution. In the context of molecular docking, this framework can be used to model distributions over protein–ligand poses. When initialized from random noise, the reverse diffusion process produces plausible binding configurations within a protein binding pocket [18, 21, 22]. Some notable examples of diffusion-based models for structure-based drug design include DiffDock [18, 21] and DiffDock-Pocket for docking, DiffSBDD [24] and MolSnapper [25] for structure-based ligand generation, and Boltz [26] and AlphaFold3 [22] for co-folding and protein structure prediction. Root-mean-square deviation (RMSD) measures the average distance between corresponding atoms of two molecular structures and is commonly used as a metric for evaluating docking accuracy. Although diffusion-based docking models such as DiffDock [18, 21] have reported strong docking performance under this metric, studies have shown that these machine-learning-based docking methods frequently generate physically implausible structures, even when the predicted poses satisfy the conventional RMSD ≤ 2 Å threshold relative to the experimentally determined binding pose [27, 28]. Similarly, diffusion-based docking models fail to consistently recover key protein–ligand interactions that can be essential for downstream lead optimization and assessing the quality of hits [29, 30]. These flaws indicate that models have not truly understood the mechanics of binding and highlight the need for approaches that explicitly incorporate criteria such as physical validity and interaction recovery in the training procedure.

We hypothesize that these issues stem from a misalignment between the standard diffusion training objective, which minimizes mean squared error on the added noise (score matching), and the downstream criteria that matter in practice, which include not only geometric accuracy as measured by RMSD but also physical plausibility and recovery of functionally important interactions. The diffusion objective optimizes physical validity and interaction recovery only indirectly, via their correlation with the training data distribution, and provides no explicit incentive to avoid physically implausible regions or preserve key interactions. For molecular docking, minimizing RMSD is not strictly equivalent to improving pose quality: while an RMSD of 0 Å would reproduce experimentally determined interactions and be physically valid (given a correct reference), reducing RMSD alone does not guarantee better interaction recovery or physical plausibility (Figure 1). For example, poses below 2 Å RMSD can still exhibit severe steric clashes or recover few interactions, indicating that geometric proximity does not uniquely determine physical validity or functional correctness.

**Figure 1.**
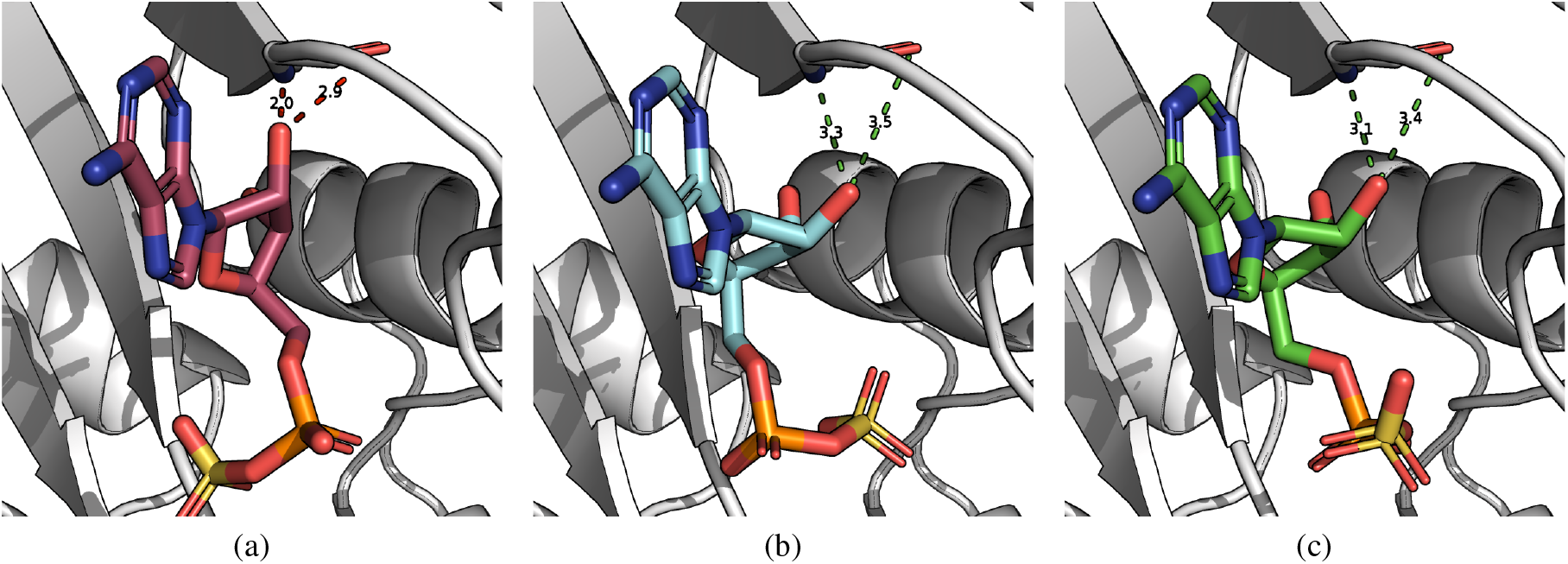
Three poses for the same protein–ligand complex are shown: (left) a sterically clashing prediction with RMSD = 1.9 Å, (center) the experimentally determined structure (PDB ID: 7HFA), and (right) a non-clashing prediction with RMSD = 2.0 Å. Despite nearly identical RMSDs, the two predictions differ in steric feasibility, illustrating that minimizing RMSD alone does not guarantee physically plausible geometries (e.g. avoiding steric clashes).

To address this gap, we introduce a reinforcement learning (RL) framework for fine-tuning diffusion-based docking models on non-differentiable downstream objectives, such as physical validity and interaction recovery. Our approach builds on recent work casting diffusion denoising as a Markov decision process that can be optimized with policy-gradient methods [31], and introduces two key innovations: early-step imitation regularization, which stabilizes training by biasing initial denoising steps toward ground-truth poses, and late-step trajectory branching, which amplifies learning signals through additional evaluations of the local refinements that often determine pose quality. We demonstrate our approach using a reward function that favors all physically plausible, interaction-preserving conformations within 2Å RMSD of the ground truth.

Applying our framework to DiffDock-Pocket [23] (yielding “DiffDock-Pocket RL”) substantially increases the fraction of near-native poses that are physically valid without post-hoc guidance or increased inference-time compute. Across all complexes, PoseBusters validity (PB-validity) increases from 58.8% to 78.1% for the top-ranked pose, and from 38.2% to 58.9% across all sampled poses. This effect is most pronounced at low sequence identity (0–30%), where the proportion of PB-valid samples increases from 24.3% to 46.4%. Despite not explicitly optimizing for this objective, the average Vina energy across generated poses improves from 2.24 kcal/mol to −2.10 kcal/mol, consistent with the model sampling more physically and energetically favorable conformations. Under stricter joint criteria, Oracle success for poses that are both ≤ 2Å RMSD to the ground truth and PB-Valid increases from 66.1% to 79.9%. DiffDock-Pocket RL outperforms the original DiffDock-Pocket and established physics-based methods on the PoseBusters benchmark set when evaluation requires physically plausible near-native poses. RL increases Top-1 success for the joint criterion ≤ 2Å RMSD & PB-valid, and yields the strongest performance when additionally requiring ≥ 50% interaction recovery. Improvements come not only from generating more physically valid poses, but also from sampling near-native poses that the baseline fails to reach. This shows that reinforcement learning increases both physical validity and the likelihood of producing accurate, near-native binding modes. When combined with physics-based local minimization and re-ranking, DiffDock-Pocket RL achieves 80.2% of Top-1 predictions with RMSD ≤ 2Å and 78.2% when physical validity is additionally required, substantially surpassing both physics-based and ML-based docking approaches on the PoseBusters set.

## 2 Methods

We propose a reinforcement learning framework for fine-tuning diffusion-based docking models using non-differentiable objectives, specifically to reward physically valid and interaction-preserving poses. Our approach builds on Deep Denoising Policy Optimization (DDPO) [31], which casts the reverse diffusion process as a Markov decision process. In this formulation, each denoising step is treated as an action, and the policy is optimized with respect to a reward defined on the final output using policy-gradient methods. This reframing allows the model to be trained directly on task-relevant criteria, such as physical validity, that are difficult to express through standard likelihood-based objectives. We introduce two key innovations over the standard DDPO framework: (1) early-step imitation regularization to stabilize training, and (2) late-step trajectory branching to provide a more informative learning signal.

### 2.1 Diffusion-Based Docking with DiffDock-Pocket

DiffDock-Pocket defines a diffusion model over the ligand pose degrees of freedom *x*_*t*_ = (tr_*t*_, rot_*t*_, tor_*t*_) ∈ ℝ^3^ × SO(3) × (*S*^1^)^*m*^, where tr_*t*_ ∈ ℝ^3^ is the ligand translation, rot_*t*_ ∈ SO(3) the rotation, and tor_*t*_ ∈ (*S*^1^)^*m*^ the *m* torsion angles, conditioned on the protein pocket representation *c*. In the forward process, each component is perturbed by independent Brownian motion with diffusion schedule *g*_*κ*_(*t*), for *κ* ∈ { tr, rot, tor }. The learned reverse process is specified by score functions *f*_*κ*_(*x*_*t*_, *c*; *θ*) in the corresponding tangent spaces.

In discrete time, the reverse transition is written in Euler–Maruyama form as Gaussian updates in the tangent coordinates for each component, where Δ*t* denotes the timestep size:

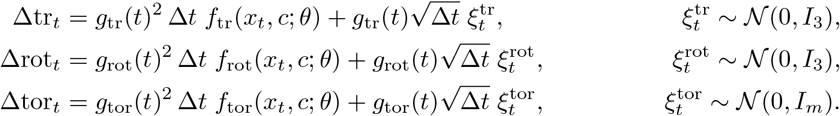

These increments are applied to update the pose as

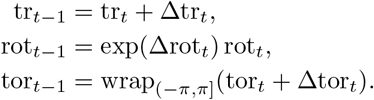

Together, these updates define a Markov transition *p*_*θ*_(*x*_*t* −1_ | *x*_*t*_, *c*), with mean drift determined by the learned scores *f*_*κ*_ and stochasticity from the Gaussian noises 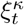. Since the reverse transition defines a stochastic distribution over pose updates, it can be interpreted as a policy. This makes it possible to optimize parameters *θ* with reinforcement learning to increase the probability of trajectories that terminate in high-reward, physically valid poses.

### 2.2 Reverse Diffusion as a Markov Decision Process

We formulate the reverse diffusion process as a Markov Decision Process (MDP), {*S, A, P, R, γ*}, over timesteps *t* = *T, T* − 1, …, 0 as follows.

#### State space (*S*)

The state at time *t* is given by *s*_*t*_ = (*x*_*t*_, *c*), where *x*_*t*_ = (tr_*t*_, rot_*t*_, tor_*t*_) represents the ligand pose at timestep *t* with translation tr_*t*_ ∈ ℝ^3^, rotation rot_*t*_ ∈ SO(3), and torsion angles tor_*t*_ ∈ (*S*^1^)^*m*^, and *c* is the conditioning protein structure.

#### Action space (*A*)

The action at time *t* is given by *a*_*t*_ = (Δtr_*t*_, Δrot_*t*_, Δtor_*t*_), where Δtr_*t*_ ∈ ℝ^3^ is the translational update, Δrot_*t*_ ∈ ℝ^3^ is an axis-angle rotation update applied via rot_*t*−1_ = exp(Δrot_*t*_)rot_*t*_, and Δtor_*t*_ ∈ ℝ^*m*^ are the *m* torsion updates, with torsion angles wrapped to (−*π, π*] after application.

#### Transition dynamics (*P*)

Given state *s*_*t*_ and (sampled) action *a*_*t*_, the next state *s*_*t*−1_ is obtained by applying the ligand pose updates from Section 2.1 deterministically, i.e. 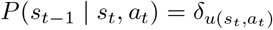, where *u* denotes the deterministic pose update and *δ* represents the Dirac delta function. All stochasticity in the MDP therefore arises from the policy, not the environment.

#### Reward (*R*)

The reward *R* is based on the final state *s*_0_ = (*x*_0_, *c*) and zero otherwise, i.e.

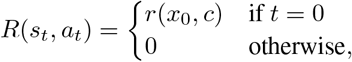

and thus the cumulative reward of each trajectory is equal to *r*(*x*_0_, *c*). Full details of the reward function are given in Section 2.6.

#### Discount factor (*γ*)

Since rewards are zero except at *t* = 0, the discounted return at timestep *t* in a trajectory is *γ*^*t*^*r*(*x*_0_, *c*) for *γ* ∈ [0, 1].

#### Policy (*π*_*θ*_)

The policy *π*_*θ*_ that determines the action *a*_*t*_ based on the current state *s*_*t*_ is parameterized by the score functions *f*_*κ*_(*x*_*t*_, *c*; *θ*) for *κ* ∈ { tr, rot, tor }. Given state *s*_*t*_, actions are sampled from Gaussian random variables for each degree of freedom, with noise sampled independently for each component:

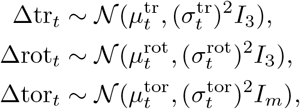

where, for each component *κ* ∈ {tr, rot, tor }, the mean and variance are determined by the score function and diffusion schedule:

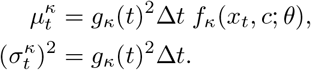

The model thus outputs the mean of the action distribution, while the stochasticity arises from sampling noise according to the diffusion schedule at each timestep. The policy likelihood factorizes across the three separate components with independent noise drawn for each:

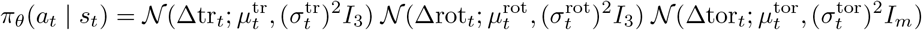

Because the policy is Gaussian with mean determined by the score functions, the log-likelihood of any action taken is tractable and differentiable with respect to *θ*, allowing direct gradient-based optimization of the RL objective.

### 2.3 Policy Gradient Objective

We optimize the policy using a policy gradient objective with importance sampling. Given a trajectory *τ* = (*s*_*T*_, *a*_*T*_, …, *s*_1_, *a*_1_, *s*_0_) sampled from the policy 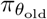, where *θ*_old_ denotes the parameters used to sample the trajectory and *θ* the current parameters that are being updated, we normalize rewards across trajectories from the same complex in the batch via z-score normalization to obtain a normalized reward 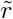. Following [31], we then optimize the importance-sampled policy-gradient objective

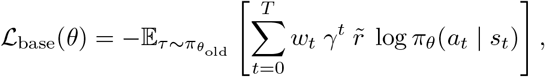

where the per-step importance ratio 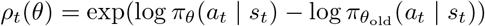 is clipped symmetrically to prevent excessively large policy updates: *w*_*t*_ = clip(*ρ*_*t*_(*θ*), 1 − *ϵ*, 1 + *ϵ*), with *ϵ >* 0.

### 2.4 Early-Step Expert Guidance

Credit assignment is a central challenge in reinforcement learning [32], where a scalar return must be attributed back to a sequence of actions that may contribute unequally to the final outcome. In long-horizon settings with only terminal rewards, standard policy-gradient methods typically propagate a trajectory-level training signal uniformly across all timesteps, which can lead to high-variance updates and weak or misaligned feedback, especially for early actions.

In diffusion-based docking, the credit-assignment problem is exacerbated by the denoising structure of the trajectory. The reward is based solely on the final pose, *x*_0_, yet the policy must choose a sequence of translational, rotational, and torsional updates over many reverse-diffusion steps. An early update that moves the ligand in a favorable direction may still receive poor feedback if later steps degrade the pose, while an early unfavorable update may be rewarded if subsequent denoising recovers a good final structure. As a result, training signals derived solely from terminal outcomes can be poorly aligned with the quality of early decisions.

To mitigate this, we introduce expert guidance for the early, high-noise portion of the trajectory, specifically timesteps *t* ≥ *T*_*E*_, where *T*_*E*_ denotes the cutoff in reverse-diffusion time (Fig. 2). Early steps are regularized towards an approximate expert action that steers the ligand towards its (known) ground-truth pose, while later steps are updated using the actions from the sampled trajectory and the terminal reward.

**Figure 2.**
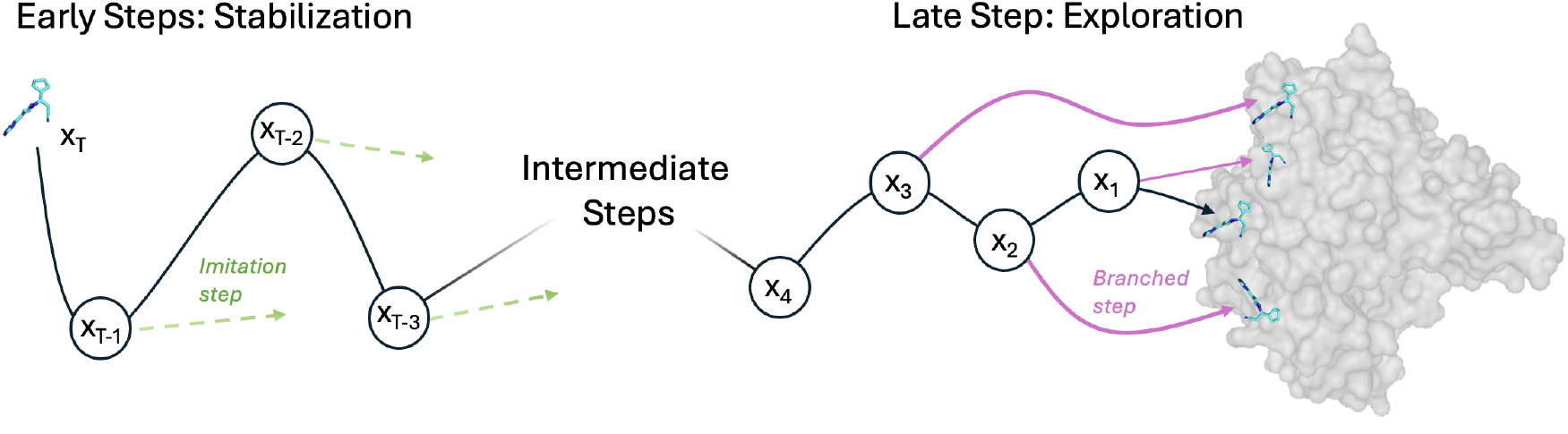
Training rollout with early imitation step guidance and late-step trajectory branching. Early denoising steps use expert actions (green arrows) that steer the ligand toward the binding pocket and ground-truth ligand. Later steps branch (pink arrows) by resampling noise to grow a binary tree of trajectories from shared intermediate states. Shared steps receive rewards averaged across all descendant leaf trajectories, while steps unique to a single leaf receive that leaf’s own reward.

For each component (translation, rotation, and torsion), we define an approximate optimal action 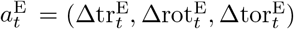 that moves the current ligand state toward its experimentally-determined ground-truth pose, scaled by the diffusion step to match the denoising schedule:

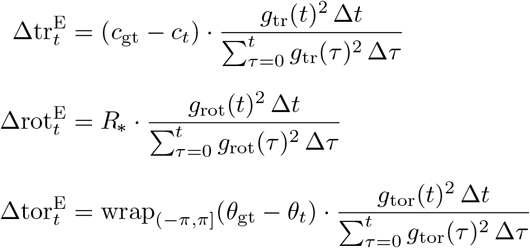

where *c*_*t*_ and *c*_gt_ are the ligand and ground-truth centers of geometry, *R*_∗_ is the optimal rotation from Kabsch alignment, and the torsion differences are wrapped to stay within (−*π, π*].

We replace the standard DDPO loss with the hybrid objective

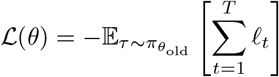

where

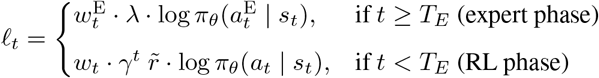

Here, *λ >* 0 is a fixed positive weighting coefficient on the expert log-likelihood term, and 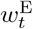 is a clipped trust region weight to prevent instability when expert actions lie far outside the current policy distribution.

This expert term is structurally equivalent to a behavior cloning loss applied only to early timesteps, in that it maximizes the likelihood of expert actions 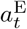 under the policy. However, the states *s*_*t*_ are still generated on-policy by 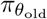, and only updates in the noisy early part of the trajectory are regularized in this way. In practice, the expert phase stabilizes credit assignment for early steps by steering trajectories toward the ground truth, while the RL phase focuses on fine-grained, reward-driven refinement of the final pose.

### 2.5 Late-Step Trajectory Branching

To obtain a denser training signal near the end of the reverse trajectory, where actions are most influential for the final pose and terminal reward, we introduce trajectory branching over a window of late denoising steps (Fig. 2). During this phase, we grow a binary tree of trajectories by resampling noise at each branching timestep from shared intermediate states. Specifically, we branch at each of the four timesteps in the window *t* ∈ { 8, …, 5 }, with each active trajectory splitting into two descendants at each branching point. This produces 16 leaf poses per complex, all sharing the same early history but diverging progressively over the branching window.

Each leaf pose *x*_0,*ℓ*_ arises from a distinct sequence of actions during and after the branching window. Because our reward *r*(*x*_0_, *c*) depends on discrete structural and interaction-level criteria, it is a highly non-linear function of the final geometry. Small local perturbations at late denoising steps can therefore induce abrupt changes in reward, for example by introducing or resolving steric clashes or disrupting key contacts. By generating a structured ensemble of end states that progressively diverge from a shared prefix, trajectory branching reveals how these non-linear criteria respond to fine-grained geometric variations at different divergence depths. This provides more informative feedback about which specific refinements turn an almost-valid pose into a valid one, or conversely push a good pose across the decision boundary into an invalid configuration. Since branches share computation up to each branching point, a single branched rollout yields 16 leaf poses without requiring 16 independent trajectory samples.

#### Reward assignment

Rewards are first normalized within each complex via z-score normalization, as described in Section 2.3. Within each rollout, we then use the tree structure to assign per-step rewards. Each internal node *v* at depth *d* (corresponding to branching timestep *t*_*d*_) is the shared ancestor of a subtree of leaves ℬ_*v*_ ⊆ ℬ, and receives the mean normalized reward of all its descendants

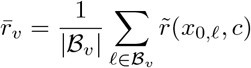

The per-step reward for an action at timestep *t* belonging to leaf trajectory *ℓ* is then

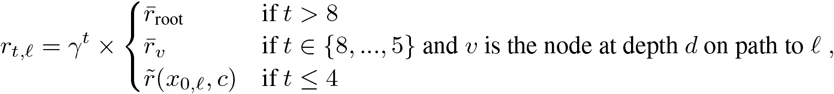

where 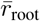 is the mean normalized reward over all 16 leaves in the rollout, and *γ* is the discount factor. Steps before branching begins receive the global rollout mean; steps at each branching timestep in {8, …, 5 } receive the mean normalized reward of their descendant subtree; and steps in the final unshared portion of each trajectory *t* ≤ 4 receive that leaf’s own normalized reward, decayed by *γ*^*t*^.

This assignment strengthens the learning signal in two ways. First, aggregating rewards across a node’s descendant leaves yields a lower-variance estimate of the quality of the policy at that state, improving credit assignment to earlier shared steps. Second, contrasting rewards across branches that differ only in their final denoising steps helps the policy resolve the sharp boundary between physically valid and invalid poses, rather than learning from a single, potentially unrepresentative outcome.

Each branching point thus contributes likelihood terms for all trajectories passing through it, concentrating gradient information at late decision points. In practice, this amplifies the number of gradient updates in late steps and allows greater exploration of small perturbations without requiring multiple independent full trajectories to be generated. We branch over the window *t* ∈ {8, …, 5 }, chosen by inspecting generated trajectories and the noise schedule, and observing that meaningful pose variation and the majority of reward-relevant geometric changes occur predominantly in the final steps of the denoising process.

### 2.6 Experimental Setup

#### Reward

We use a terminal reward *r*(*x*_0_, *c*) that rewards both physical validity and predictions that are close to the ground truth. Specifically, we assess the final pose using the PoseBusters physical validity checks and assess whether the generated pose is within 2Å RMSD of the ground truth structure. The reward is the proportion of all checks passed by the final pose. We use a discount factor of *γ* = 0.95. While the PoseBusters package includes a check for whether the RMSD between the generated pose is ≤ 2Å from the ground truth, we define a structure to be PoseBusters valid (PB-valid) if it passes all of the PoseBusters plausibility checks excluding the RMSD ≤ 2Å check (i.e. all intermolecular, intramolecular, and chemical validity checks that do not require knowledge of the ground-truth pose). Full details of the checks and thresholds are provided in the PoseBusters paper [27].

#### Dataset

To train our model, we use the PDBbind 2020 refined set with the same train-validation splits that were used to train DiffDock-Pocket [23] in order to ensure that any improvement is as a result of our modified training procedure as opposed to a change in data. Complexes whose ground-truth poses fail PB-validity checks are removed, resulting in the exclusion of 562 out of 4503 complexes (12.5%), thereby ensuring clean training signals. Throughout our analysis, we use the PoseBusters benchmark set to assess model performance. This benchmark comprises 308 high-quality protein–ligand complex structures. All entries were deposited in the Protein Data Bank (PDB) after the structures used for training, and sequence similarity metrics are included to assess model generalization.

#### Evaluation metrics

We evaluate pose quality using four criteria that are generally reported cumulatively as increasingly stringent success definitions: (i) coordinate accuracy, defined as ligand RMSD ≤ 2Å to the experimental reference; (ii) physical plausibility, defined as passing all PB-validity checks; (iii) interaction recovery (IR) of at least 50%; and (iv) IR of at least 75%. Interaction recovery is calculated using ProLIF as described in [29], considering hydrogen bonds, halogen bonds, *π*-stacking, cation-*π*, and ionic interactions. Success rates are primarily reported by applying these criteria sequentially. To assess generalizability, we stratify results by sequence identity between the target protein and the most similar protein in the training set, using 0–30%, 30–95%, and 95–100% sequence identity ranges, in line with [27]. We run DiffDock-Pocket and the fine-tuned DiffDock-Pocket RL for three independent inference runs and report results as mean ± standard deviation across runs. Statistical significance of differences between models is assessed using McNemar’s test on per-complex binary success outcomes.

#### Pose ranking and Top-1 vs Oracle

Many methods generate multiple candidate poses per complex and use a separate ranking step to select a final prediction. DiffDock-Pocket follows this paradigm, assigning each candidate a score using a separate confidence model. For each complex, we generate 40 candidate poses per method. We report performance using two evaluation protocols: “Top-1”, the highest-ranked pose under the fixed confidence model, and “Oracle”, the best pose among all sampled candidates for that complex according to the evaluation criterion. Thus, Top-1 reflects end-to-end performance including pose selection, while Oracle isolates improvements in the generative model independent of the confidence model. Note we do not modify the confidence model during training.

#### Comparison to other models

We compare against AutoDock Vina [10], GOLD [12], and the original DiffDock-Pocket model [23]. We include Vina and GOLD because physics-inspired docking methods remain a gold standard for generating physically valid poses, providing a strong reference point for plausibility-oriented evaluation. We include the baseline DiffDock-Pocket model to isolate the effect of our reinforcement-learning fine-tuning procedure relative to the same diffusion architecture and inference pipeline. For broader context, we also report the performance for a set of representative ML-based docking methods, namely DiffDock [18], EquiBind [14], TankBind [15], UniMol [33], and DeepDock [34]. We use the same poses for these methods that were used in the original PoseBusters analysis [27]. Additional implementation details are provided in the PoseBusters paper [27].

## 3 Results

### 3.1 Physically Motivated Rewards Improve Pose Validity and Plausibility

Our model, trained with RL using the proportion of PoseBusters validity checks passed as a reward, results in a substantial increase in the overall proportion of valid poses generated. Across all complexes, PB-validity increases from 58.8% to 78.1% for the top-ranked pose, and from 38.2% to 58.9% across all sampled poses. A breakdown of performance for each of the PoseBusters criteria for the baseline and RL-fine-tuned models is provided in Appendices B and C. Hereafter, we refer to the RL fine-tuned model as “RL” and the original DiffDock-Pocket model as “Baseline”. Figure 3 shows the proportion of PB-valid poses for the baseline DiffDock-Pocket model and our RL-fine-tuned model across sequence identity ranges. RL increases the overall PB-valid fraction across all sequence identity ranges, with the most substantial improvements for the most out-of-distribution targets with the lowest similarity to training proteins. For targets with 0–30% sequence identity to the training set, overall PB-validity increases from 24.3% to 46.4%. For targets closer to the training distribution, RL training also yields substantial improvements, increasing overall PB-validity from 36.4% to 56.8% for targets with 30–95% sequence identity and from 51.5% to 71.2% for targets with 95–100% sequence identity. These results indicate that directly optimizing PB-validity reliably improves physical plausibility across both in- and out-of-distribution settings.

**Figure 3.**
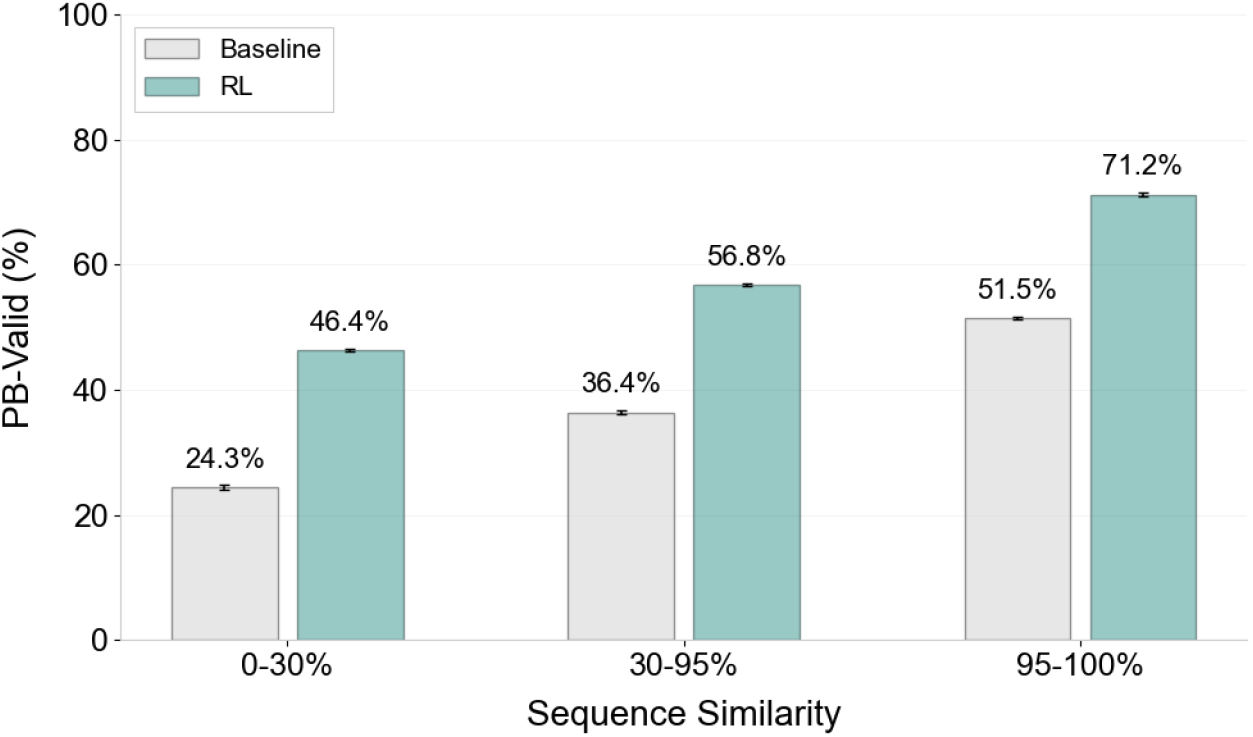
PB-validity of sampled poses. Fine-tuning DiffDock-Pocket with reinforcement learning (“RL”) significantly increased the proportion of physically-valid poses across all sequence identity ranges between targets and the most similar training example compared to the original DiffDock-Pocket model (“Baseline”).

To evaluate whether our RL-fine-tuned model has improved the quality of docked poses as defined by a physics-based scoring function, we compute Vina energies for every pose generated by both the baseline DiffDock-Pocket model and our RL-trained model. Figure 4 shows the resulting energy distributions. While Vina energies are an imperfect proxy for binding quality, high energies typically indicate implausible poses, for example due to severe steric clashes, while lower energies suggest greater physical plausibility. Poses generated by the RL model show a substantial shift toward lower energies, decreasing the mean Vina score from 2.24 kcal/mol to -2.10 kcal/mol, an improvement of 4.34 kcal/mol. This indicates that optimizing for PB-validity both increases physical plausibility directly and also shifts the distribution of generated poses toward lower Vina energies.

**Figure 4.**
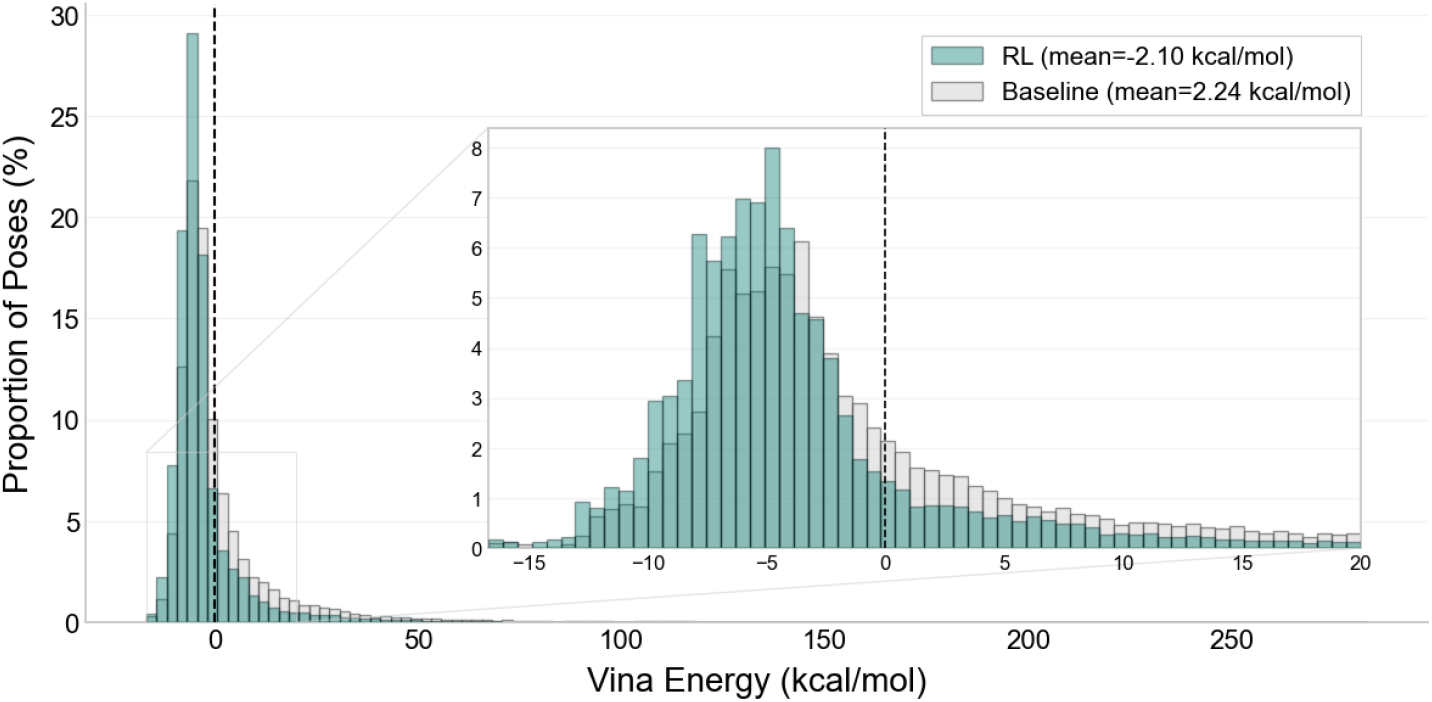
Distribution of Vina energies. Histogram of predicted binding energies (kcal/mol) for poses generated by the baseline model and the RL-optimized model. Reinforcement learning produces a substantial improvement in docking score, decreasing the average Vina energy from 2.24 kcal/mol to -2.10 kcal/mol.

### 3.2 Reinforcement Learning Improves Performance Under All Evaluation Criteria

Reinforcement learning improves performance across all evaluation criteria, with the most notable gains occurring when physical validity is required (Table 1). Top-1 performance increases from 46.2% to 58.8% when requiring ≤ 2Å RMSD and PB-valid, and from 38.9% to 49.7% when additionally requiring at least 50% interaction recovery. Substantial improvements are also observed for the Oracle evaluation, which measures whether any generated pose satisfies the criteria regardless of ranking, indicating that RL more reliably generates at least one pose meeting strict combined criteria across complexes. We additionally report results stratified by sequence similarity to the training set (Appendix G), showing consistent improvements over the baseline across similarity bins, with the greatest improvements on low-similarity targets (0–30%).

**Table 1.** Success rates under increasingly strict quality criteria on the PoseBusters benchmark set (n=308). Our RL fine-tuned version of DiffDock-Pocket (“RL”) improves performance for all criteria compared to the base DiffDock-Pocket model (“Baseline”).

| Metric | Top-1 (%) |  | Oracle (%) |  |
| --- | --- | --- | --- | --- |
|  | Baseline | RL | Baseline | RL |
| RMSD $\leq 2\text{\AA}$ | 66.8 ± 0.5 | **69.0 ± 2.8<sup>ns</sup>** | 78.0 ± 1.4 | **84.8 ± 0.4 \*\*\*\*** |
| & PB-valid | 46.2 ± 1.0 | **58.8 ± 1.7 \*\*\*\*** | 66.1 ± 0.7 | **79.9 ± 0.3 \*\*\*\*** |
| & $\geq 50\%$ IR | 38.9 ± 0.5 | **49.7 ± 1.7 \*\*\*\*** | 61.4 ± 0.9 | **74.7 ± 0.6 \*\*\*\*** |
| & $\geq 75\%$ IR | 28.5 ± 0.8 | **35.8 ± 1.8 \*\*\*\*** | 50.9 ± 1.0 | **64.8 ± 1.0 \*\*\*\*** |

Values are reported as mean ± standard deviation across runs. Significance assessed using McNemar’s test on per-complex binary success outcomes, pooled across run pairs. ns: not significant ( $p \geq 0.05$ ); \*\*\*\*: $p < 0.0001$ .

**Table 2.**
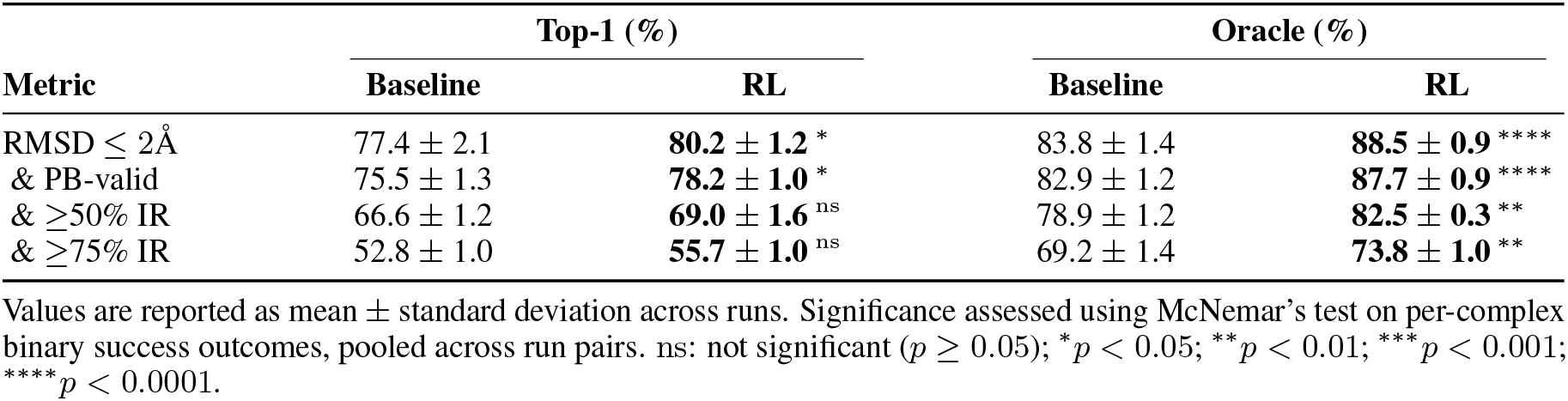
Success rates under increasingly strict quality criteria on the PoseBusters benchmark set (n=308) after Vina minimization and GNINA re-ranking. Our RL fine-tuned model (“RL”) outperforms the base DiffDock-Pocket model (“Baseline”) across all metrics, showing that improvements persist after post-hoc minimization.

| Metric | Top-1 (%) |  | Oracle (%) |  |
| --- | --- | --- | --- | --- |
|  | Baseline | RL | Baseline | RL |
| RMSD $\leq 2\text{\AA}$ | 77.4 ± 2.1 | **80.2 ± 1.2 \*** | 83.8 ± 1.4 | **88.5 ± 0.9 \*\*\*\*** |
| & PB-valid | 75.5 ± 1.3 | **78.2 ± 1.0 \*** | 82.9 ± 1.2 | **87.7 ± 0.9 \*\*\*\*** |
| & $\geq 50\%$ IR | 66.6 ± 1.2 | **69.0 ± 1.6 <sup>ns</sup>** | 78.9 ± 1.2 | **82.5 ± 0.3 \*\*** |
| & $\geq 75\%$ IR | 52.8 ± 1.0 | **55.7 ± 1.0 <sup>ns</sup>** | 69.2 ± 1.4 | **73.8 ± 1.0 \*\*** |

Values are reported as mean ± standard deviation across runs. Significance assessed using McNemar’s test on per-complex binary success outcomes, pooled across run pairs. ns: not significant ( $p \geq 0.05$ ); \* $p < 0.05$ ; \*\* $p < 0.01$ ; \*\*\*\* $p < 0.001$ ; \*\*\*\* $p < 0.0001$ .

Although our model is trained using only PB-validity as the reward, we also observe improvements in interaction recovery. A plausible explanation is that reducing steric clashes places poses in more plausible contact geometries with the protein, and the increased diversity of valid poses improves the chance of recovering native contacts, which is consistent with our analysis in Section 3.5. The cumulative structure of our evaluation criteria also contributes, as interaction recovery is assessed jointly with physical validity and geometric accuracy, so increases in the number of PB-validity poses increase the pool of poses eligible to satisfy stricter criteria.

### 3.3 Reinforcement Learning Maintains Improvement After Physics-Based Relaxation

Previous work has shown that post-hoc energy minimization can recover some performance lost due to physical implausibility [27]. To investigate the effect of post-hoc minimization, we minimized all generated poses using smina [35] with the Vina scoring function and re-ranked poses using GNINA [13], and report Top-1 and Oracle metrics. RL outperforms the baseline across both Top-1 and Oracle metrics, with statistically significant improvements for RMSD ≤ 2Å and the combined criterion of RMSD ≤ 2Å and PB-valid. For the criteria additionally requiring interaction recovery, Top-1 differences are not statistically significant, though Oracle improvements remain significant, suggesting that while re-ranking does not consistently identify them, minimized RL poses more reliably contain at least one pose meeting strict combined criteria across complexes.

Although minimization narrows the gap between models by improving baseline performance relatively more than RL (consistent with the baseline having greater room for improvement due to lower initial physical validity), the RL model maintains an advantage across all Oracle metrics after minimization.

### 3.4 DiffDock-Pocket RL Outperforms Physics-Based and Machine Learning Docking Methods

Table 3 compares Top-1 performance for physics-based docking methods and machine learning-based approaches on the PoseBusters benchmark. Prior to physics-based refinement, DiffDock-Pocket RL is already competitive with established physics-based docking methods and outperforms almost all ML-based approaches. DiffDock-Pocket RL achieves 69.0% of predictions with RMSD ≤ 2Å, outperforming Vina (59.7%) and GOLD (58.1%). When physical validity is additionally required, DiffDock-Pocket RL achieves 58.8% for RMSD ≤ 2Å and PB-valid, again exceeding Vina (58.1%) and GOLD (54.2%). Under stricter interaction recovery criteria, DiffDock-Pocket RL attains 49.7% for ≥ 50% IR compared with 44.8% for Vina and 45.5% for GOLD. For the most stringent criterion at ≥ 75% IR, DiffDock-Pocket RL reaches 35.8%, approaching the best physics-based performance of 37.7% achieved by GOLD.

**Table 3.** Top-1 success rates (%) under increasingly strict quality criteria on the PoseBusters benchmark set (n=308) comparing physics-based and machine learning-based docking methods.

| Method | RMSD $\leq 2$ Å (%) | & PB-valid (%) | & $\geq 50\%$ IR (%) | & $\geq 75\%$ IR (%) |
| --- | --- | --- | --- | --- |
| **Physics-Based** |  |  |  |  |
| Vina [10] | 59.7 | 58.1 | 44.8 | 35.1 |
| GOLD [12] | 58.1 | 54.2 | 45.5 | 37.7 |
| **Machine Learning-Based** |  |  |  |  |
| DeepDock [34] | 19.5 | 5.2 | 3.2 | 2.3 |
| DiffDock [18] | 38.0 | 12.3 | 9.4 | 7.1 |
| EquiBind [14] | 1.9 | 0.0 | 0.0 | 0.0 |
| TankBind [15] | 15.9 | 3.9 | 1.9 | 1.3 |
| UniMol [33] | 20.8 | 1.9 | 0.6 | 0.6 |
| DiffDock-Pocket [23] | 66.8 | 46.2 | 38.9 | 28.5 |
| GNINA* [13] | 73.7 | 72.4 | 56.2 | 40.9 |
| DiffDock-Pocket RL | 69.0 | 58.8 | 49.7 | 35.8 |
| DiffDock-Pocket RL++ | **80.2** | **78.2** | **69.0** | **55.7** |

\*GNINA combines physics-based search with a machine learning-based scoring function to rank poses.

As shown in Section 3.3, the performance of DiffDock-Pocket RL can be improved by minimizing poses using Vina and re-ranking using GNINA at minimal additional computational cost, in part by replacing the original DiffDock-Pocket confidence model. This approach, denoted “DiffDock-Pocket RL++”, achieves the best performance of all methods tested, with 80.2% of predictions with RMSD ≤ 2Å and 78.2% when PB-valid is additionally required. Under stricter interaction recovery thresholds, performance reaches 69.0% for ≥ 50% IR and 55.7% for ≥ 75% IR. These results surpass both physics-based and machine learning-based docking approaches with comparable binding site definitions. The improvement over other ML methods is particularly substantial when physical validity is required, with DiffDock-Pocket RL++ achieving 78.2% for RMSD ≤ 2Å and PB-valid, compared with 72.4% for GNINA, 46.2% for DiffDock-Pocket, 12.3% for DiffDock, and below 10% for all other ML baselines. Overall, this demonstrates that reinforcement learning can effectively enhance diffusion-based docking models, and that combining learned generative models with physics-based refinement yields strong docking performance. For completeness, we report results for machine learning docking methods with their corresponding pocket sizes in Appendix A.

### 3.5 Analysis of Generated Poses

In this section, we analyze the poses generated by DiffDock-Pocket RL and the original DiffDock-Pocket model to understand the reason for the observed performance differences. This analysis reveals how RL training changes the distribution of generated poses and identifies the failure modes it corrects.

Figure 5 shows the distribution of average pairwise RMSD between all valid poses for each complex, examining whether the RL model generates diverse physically valid poses or reproduces the same valid pose many times for a given complex. RL exhibits higher mean pairwise RMSD (2.21Å) relative to the baseline DiffDock-Pocket model (1.25Å) alongside more complexes with PB-valid poses (291 vs 244). The simultaneous increase in diversity and validity, alongside the disproportionate increase in physically valid poses observed for low-homology targets, suggests the model has learned generalizable principles of validity rather than relying on memorization of valid training poses.

**Figure 5.**
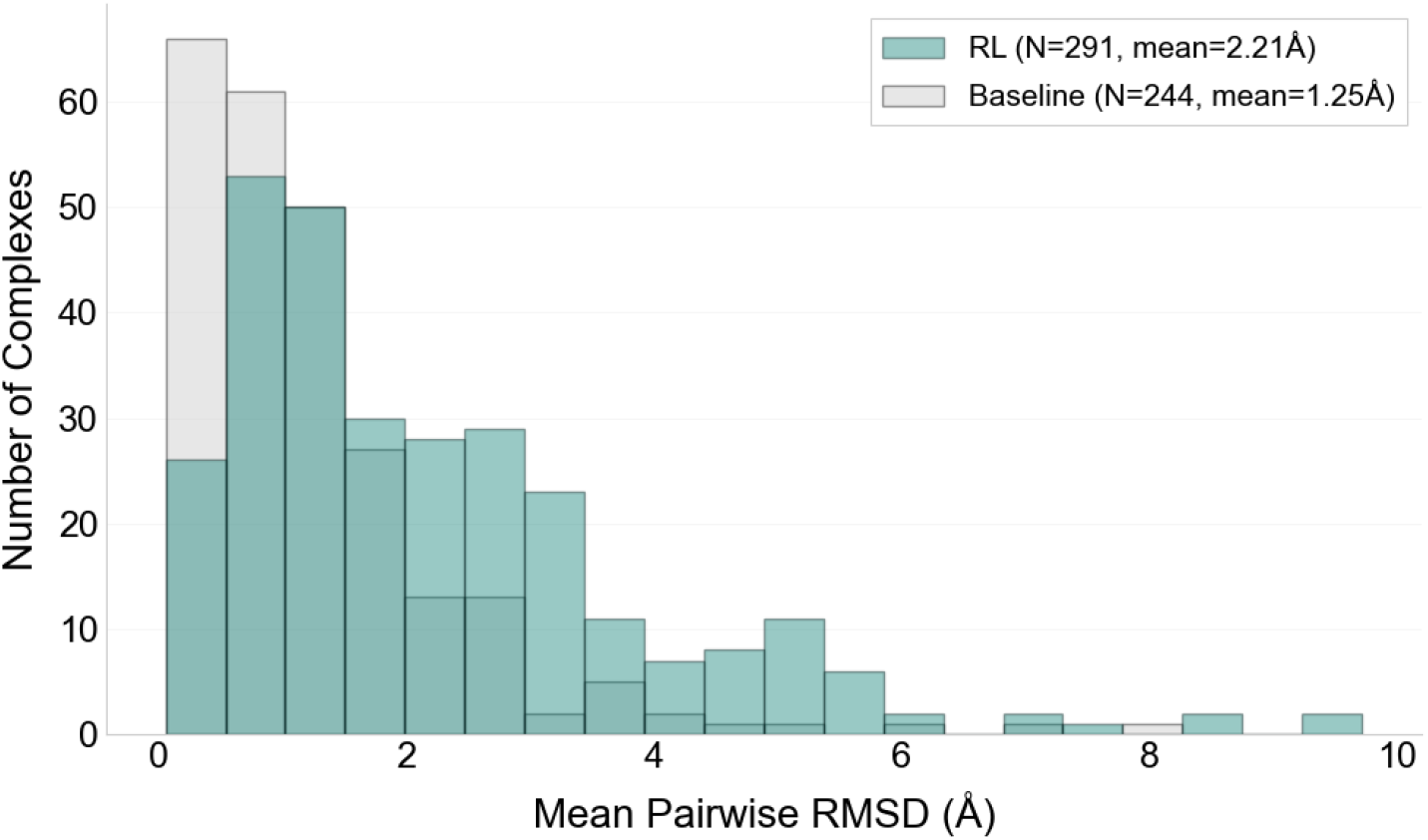
Diversity of PB-valid poses. Distribution of average pairwise RMSD between PB-valid poses within the full generated set for each of the PoseBusters benchmark complexes (N=308). The RL model generates more diverse valid poses (mean pairwise RMSD 2.21 Å) than the baseline (1.25 Å).

We analyze the failure modes of the original DiffDock-Pocket (“Baseline”) on cases where RL succeeded and summarize the results in Table 4. We ran each model three times, generating 40 poses per complex per run. For each of the three runs, we identify complexes for which RL produced at least one pose that is both PB-valid and ≤ 2Å RMSD, while the baseline produced no such pose across all 40 generated poses. This yields 139 such cases across the three runs, corresponding to 69 unique complexes. For these cases, we classify the failure modes of the baseline model according to whether it produced PB-invalid poses with ≤ 2Å RMSD, PB-valid poses with *>*2Å RMSD, PB-invalid poses with *>*2Å RMSD, or both ≤ 2Å RMSD poses and PB-valid poses but never simultaneously. We also report the best RMSD achieved across all baseline poses and, if any were generated, the best RMSD among PB-valid baseline poses. Appendix H also provides Venn diagrams summarizing the overlap in success sets between the original DiffDock-Pocket baseline and the RL-trained DiffDock-Pocket model, stratified by sequence similarity.

**Table 4.** Failure mode analysis of the baseline DiffDock-Pocket model on cases where RL succeeds (69 unique complexes, 139 complex-run pairs). The majority of baseline failures occur because near-native poses are not PB-valid (38.1%), though RL also succeeds in many cases where the baseline fails to reach the native binding mode entirely.

| Failure mode | Count (%) | Best RMSD overall | Best RMSD among valid |
| --- | --- | --- | --- |
| $\leq 2\text{\AA}$ RMSD and Not PB-Valid | 53 (38.1%) | $1.22 \pm 0.46\text{\AA}$ | N/A |
| $> 2\text{\AA}$ RMSD and PB-Valid | 42 (30.2%) | $3.57 \pm 1.86\text{\AA}$ | $4.71 \pm 2.48\text{\AA}$ |
| $> 2\text{\AA}$ RMSD and Not PB-Valid | 22 (15.8%) | $4.07 \pm 1.88\text{\AA}$ | N/A |
| $\leq 2\text{\AA}$ RMSD or PB-Valid (Separately) | 22 (15.8%) | $1.28 \pm 0.37\text{\AA}$ | $5.54 \pm 3.74\text{\AA}$ |

Table 4 shows that, in most cases where the baseline model fails to produce a pose that is both PB-valid and ≤ 2Å RMSD, the failure occurs because near-native poses are not PB-valid. In 38.1% of cases, the baseline achieves ≤ 2Å RMSD but no PB-valid poses are generated. However, RL also succeeds in many cases where the baseline does not reach the native binding mode. In 30.2% of cases, the baseline produces PB-valid poses but none ≤ 2Å RMSD, with a mean best RMSD among valid poses of 4.71Å, suggesting that it samples physically plausible poses with an alternative binding mode. Furthermore, the mean best RMSD achieved across all poses for these cases is 3.57Å, indicating that the baseline is not close to the native binding mode even among its invalid poses. Finally, 15.8% of cases contain no PB-valid poses and no poses ≤ 2Å RMSD across all sampled poses, and a further 15.8% of cases contain both a PB-valid pose and a ≤ 2Å pose but never simultaneously, indicating that the baseline is often unable to generate poses that are both near native and valid where RL does.

## 4 Conclusion

In this work, we demonstrate that reinforcement learning can teach diffusion-based docking models to generate a higher proportion of physically valid molecular poses without sacrificing structural accuracy or increasing inference-time compute. Fine-tuning DiffDock-Pocket using a non-differentiable terminal reward based on PB-validity improves Top-1 success for poses that are both ≤ 2Å RMSD and PB-valid from 46.2% to 58.8%, and Oracle success from 66.1% to 79.9%. The most substantial improvements are observed in the lowest-homology targets (0–30% sequence identity), suggesting that the RL-based training has improved generalization.

Our primary aim was to improve the quality of the generated poses, and thus in this work we did not seek to improve or retrain the ranking model used by DiffDock-Pocket, despite the shift in pose distribution as a result of RL training. As a consequence, the improvement in Oracle performance was larger than Top-1, suggesting limitations in the ranking approach adopted. We show that minimizing poses generated by our approach using a physics-based method and re-ranking can yield substantial improvements, and recommend this approach be used in practice. DiffDock-Pocket RL++ achieves 80.2% Top-1 success for RMSD ≤ 2Å and 78.2% when physical validity is additionally required, surpassing all physics-based and ML-based docking methods tested on the PoseBusters benchmark. Improving the confidence model is a promising direction for future work and could result in more of the improvement in Oracle performance being translated into Top-1 success.

While recent work has begun to explore guidance-based approaches to improve pose validity in other applications of diffusion models in SBDD, such as Boltz-steering [26], these methods require additional forward passes and validity checks during denoising, increasing memory use and inference time. More fundamentally, guidance reshapes sampling at inference time without changing the model, so it is not expected to lead to improvements when the base model places the ligand in the wrong region of the target. Our reinforcement learning framework instead updates the learned score function so that the model itself assigns higher probability to physically valid poses and to poses that are both valid and near-native. Although our method yields substantial improvements over DiffDock-Pocket in the low sequence similarity regime, performance remains lower on the most dissimilar targets than on those closer to the training distribution. Closing the gap on the most out of distribution targets therefore remains an open challenge.

Recent cofolding tools, such as AlphaFold3 and Boltz [22, 26], and diffusion-based molecular generation models, such as DiffSBDD [24], have been shown to suffer similar failure modes, such as generating physically invalid structures [28], suggesting both could benefit from RL fine-tuning. By using non-differentiable physical criteria as training signals, we can optimize for constraints that are difficult to encode through supervised losses alone. As the field continues to develop increasingly powerful generative models for biomolecular structures, ensuring their outputs are not just accurate but also physically valid and useful for downstream screening and lead optimization will be essential for translating these advances into practical tools for drug discovery and protein engineering.

We show that diffusion-based docking models can be trained to more explicitly respect physical principles that are not captured by the standard supervised objective. Reinforcement learning enables these constraints to be incorporated directly into training through non-differentiable objectives while maintaining structural accuracy. This provides a general approach for improving the physical validity of predictions made by diffusion-based models for structure prediction and can be extended to other structure-based prediction tasks.

## Code Availability

The source code for this project is available at https://github.com/oxpig/RLDiff.

## Acknowledgments

James Broster acknowledges funding from the Engineering and Physical Sciences Research Council (EPSRC) under grant number EP/S024093/1 and from the Cambridge Crystallographic Data Centre through the SABS:R^3^ doctoral training programme. The authors acknowledge the use of resources provided by the Isambard-AI National AI Research Resource (AIRR). Isambard-AI is operated by the University of Bristol and is funded by the UK Government’s Department for Science, Innovation and Technology (DSIT) via UK Research and Innovation, and the Science and Technology Facilities Council [ST/AIRR/I-A-I/1023].

## A Pocket definitions across docking methods

The binding sites into which ligands are docked are defined as follows:

**AutoDock Vina [10]:** The search space is defined as a cubic bounding box of side length 25 Å centred at the ligand centroid.

**GOLD [12]:** The binding site is defined as a sphere of radius 25 Å centred at the ligand centroid.

**DiffDock-Pocket [23]:** Receptor C*α* atoms within 5 Å of ligand heavy atoms are selected and their mean coordinate defines the pocket centre. The pocket radius is set as the maximum distance from this centre to any ligand heavy atom plus a 10 Å buffer.

**SigmaDock [36]:** All receptor residues within a distance threshold *d*_0_ of any ligand heavy atom are included. We report *d*_0_ = 7 Å for greatest comparability with other approaches.

**UniMol [33]:** All receptor residues with any atom within 8 Å of a ligand heavy atom are included.

**DeepDock [34]:** The binding region is defined as a surface patch extending approximately 10 Å from the ligand.

**Figure 6.**
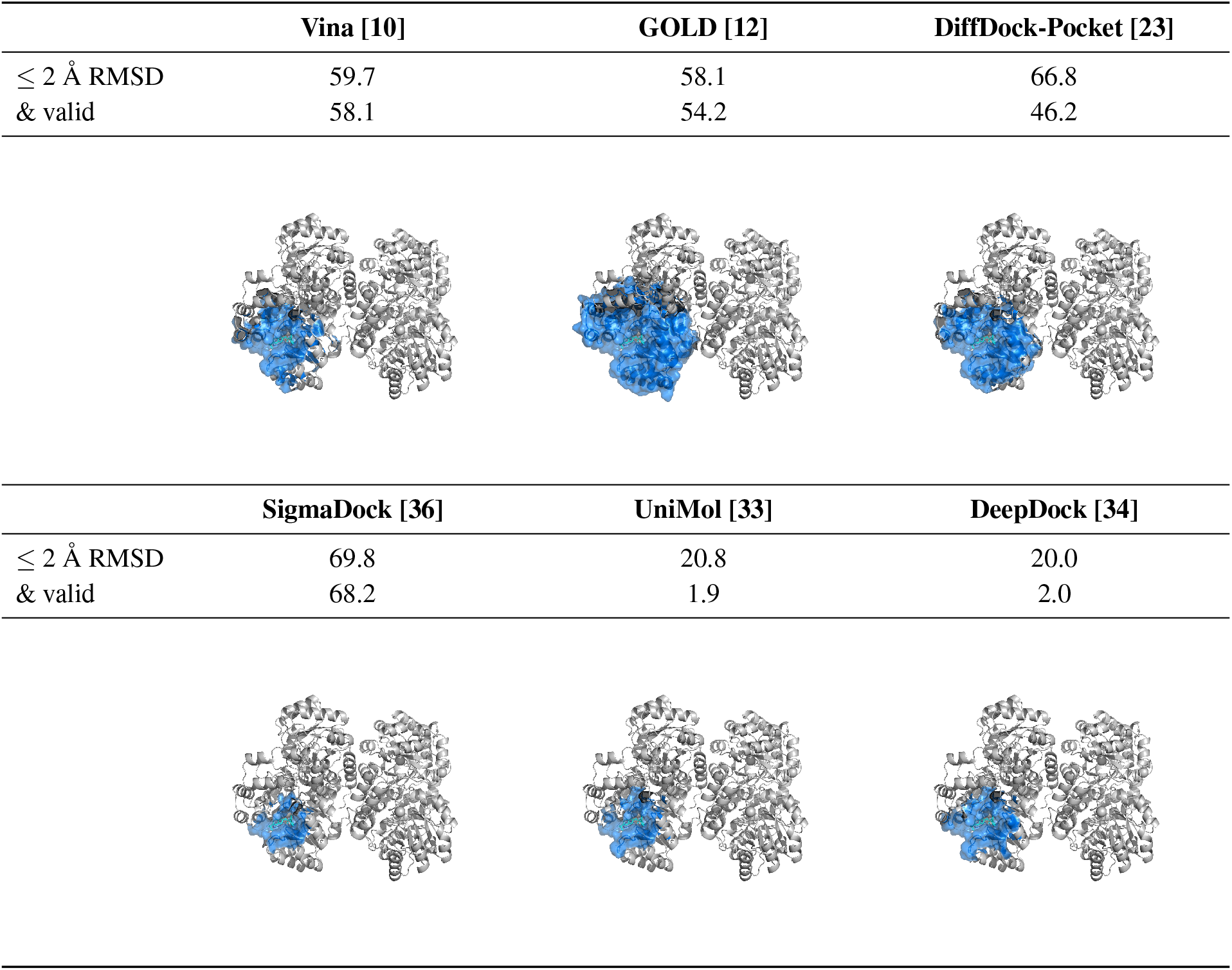
Pocket definitions and performance across docking methods. The figure shows performance alongside visualisations of the corresponding pocket definitions.

## B PoseBusters Physical Validity Checks

Figures 7 and 8 provide a breakdown of pass rates for each individual PoseBusters validity test. Figure 7 shows results for the top-ranked pose per complex, while Figure 8 aggregates across all 40 sampled poses. The largest improvements from RL training occur in the “Minimum Distance To Protein” test, improving from 60.4% to 80.3% for top-ranked poses. “Volume Overlap With Protein” also improves from 95.5% to 99.2%, indicating that RL substantially reduces steric clashes. Improvements are also observed for interaction geometry tests involving cofactors and water molecules.

**Figure 7.**
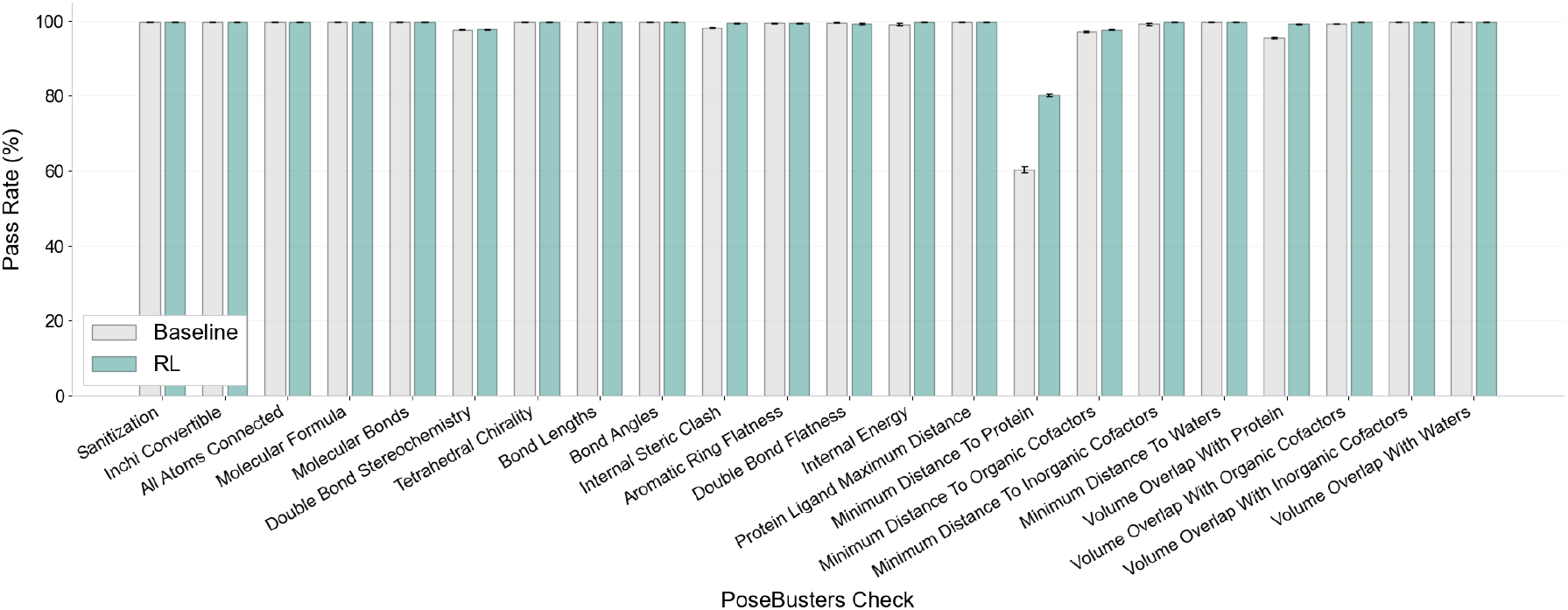
Pass rates for individual PoseBusters validity tests on top-ranked poses. Baseline model (gray) compared against RL-trained model (blue). Error bars show standard error across three independent runs.

**Figure 8.**
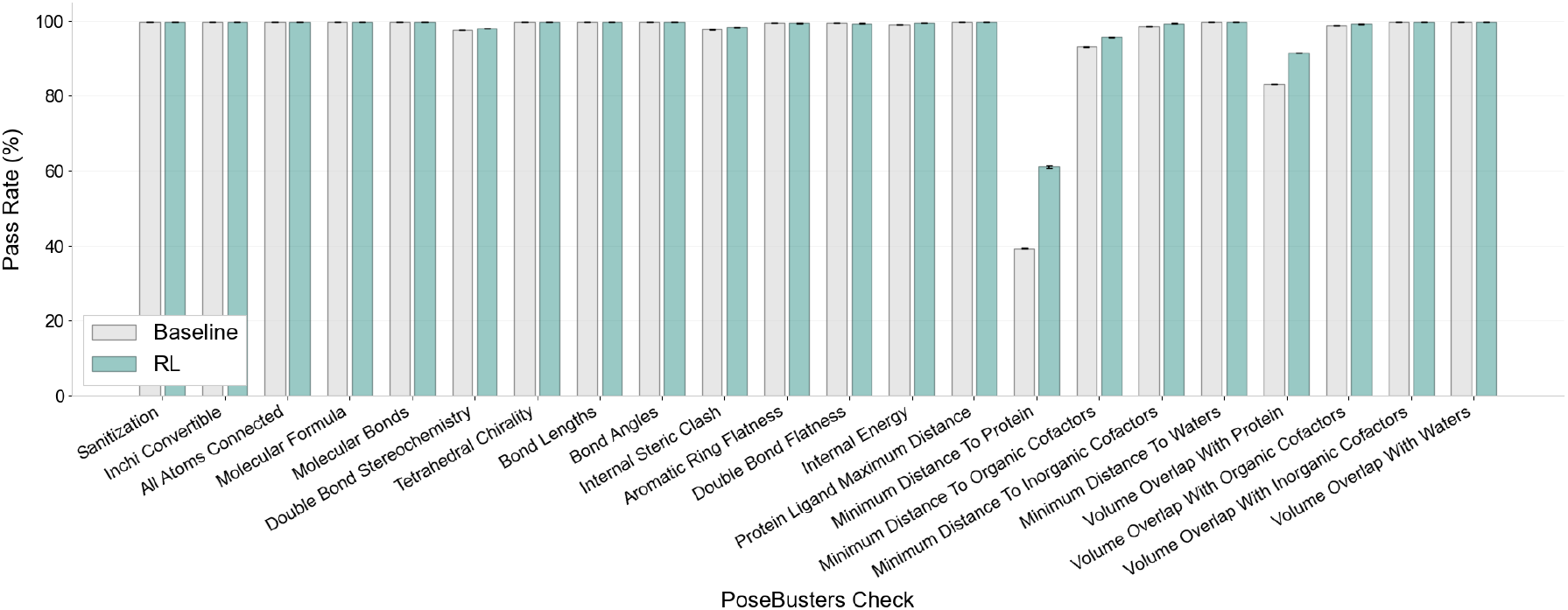
Pass rates for individual PoseBusters validity tests aggregated across all 40 sampled poses per complex. Baseline model (gray) compared against RL-trained model (blue). Error bars show standard error across three independent runs.

## C PoseBusters Validity Checks Stratified by Sequence Identity

Figures 9–11 show pass rates for individual PoseBusters validity tests stratified by sequence identity to the training set, aggregated across all poses. RL training provides the largest improvements for low-homology targets (0–30% sequence identity), where “Minimum Distance To Protein” improves from 25.3% to 48.8% (a 92.9% relative improvement), and “Volume Overlap With Protein” improves from 74.4% to 86.8%. For high-homology targets (95–100%), RL training further improves “Minimum Distance To Protein” from 53.0% to 73.5%, demonstrating consistent benefits across different sequence similarities.

**Figure 9.**
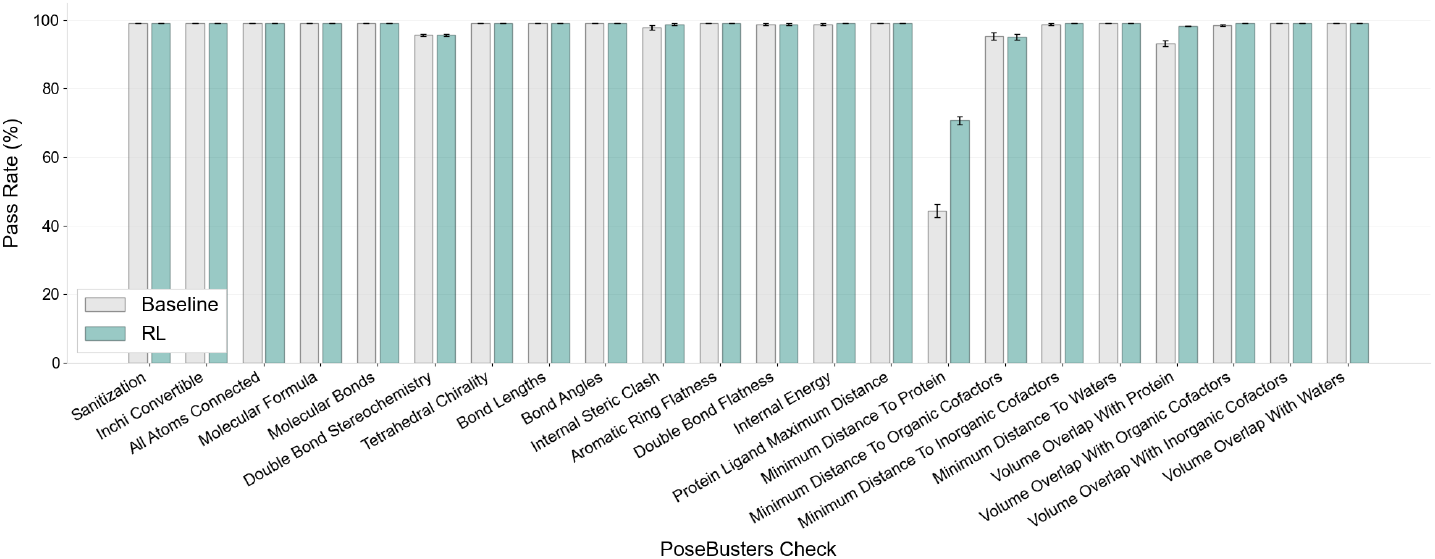
Success rates for PoseBusters checks (0-30% sequence identity).

**Figure 10.**
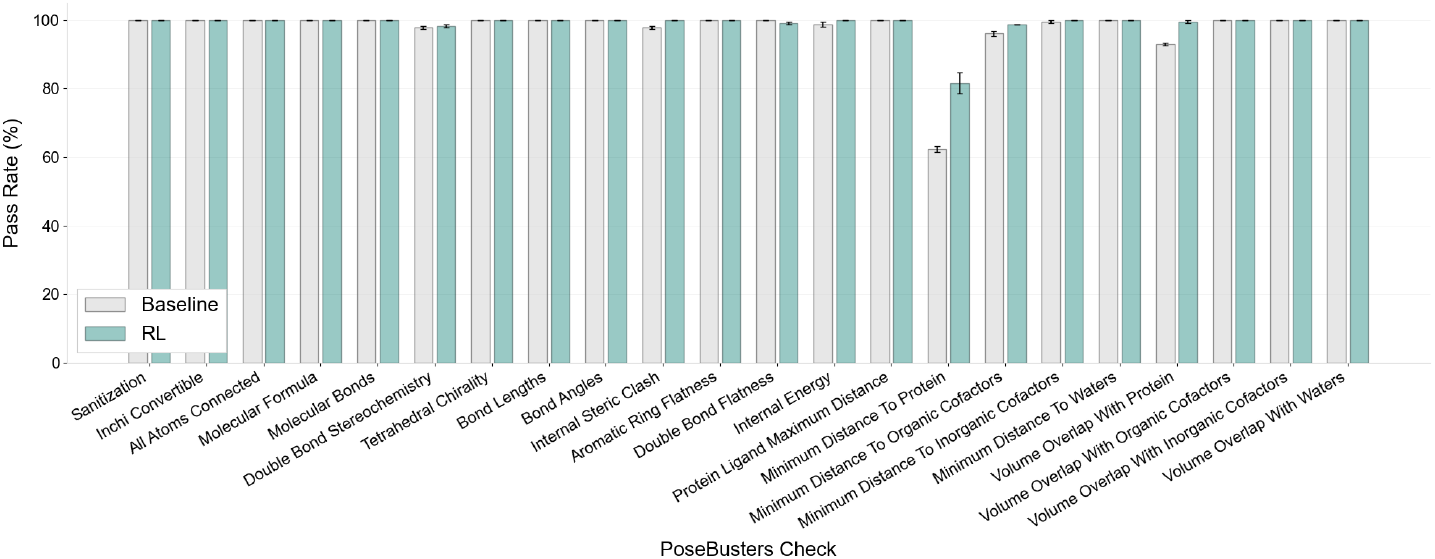
Success rates for PoseBusters checks (30-95% sequence identity).

**Figure 11.**
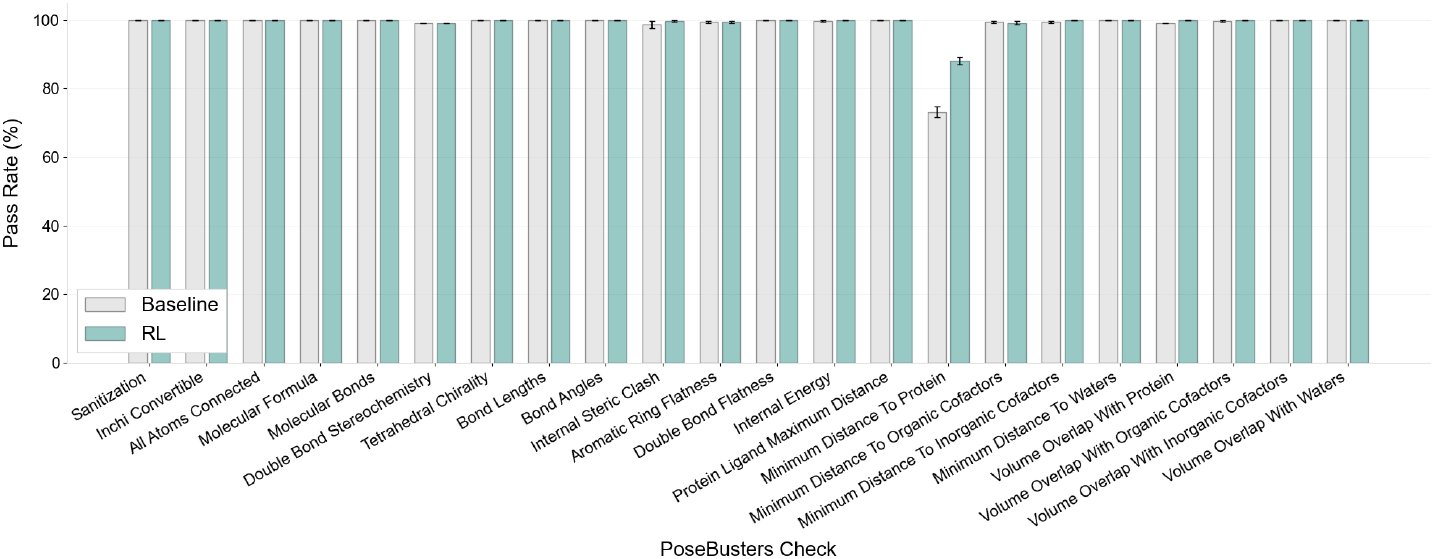
Success rates for PoseBusters checks (95-100% sequence identity).

## D Sample Efficiency

Figure 12 shows success rates for the joint criterion RMSD ≤ 2 Å and PB-valid as a function of the number of poses sampled per complex, for both Top-1 and Oracle evaluation. Under Top-1, RL consistently outperforms the baseline across all sample sizes. Under Oracle evaluation, the gap between RL and the baseline widens substantially with more samples, indicating that RL generates a richer ensemble of valid poses and benefits more from additional sampling.

**Figure 12.**
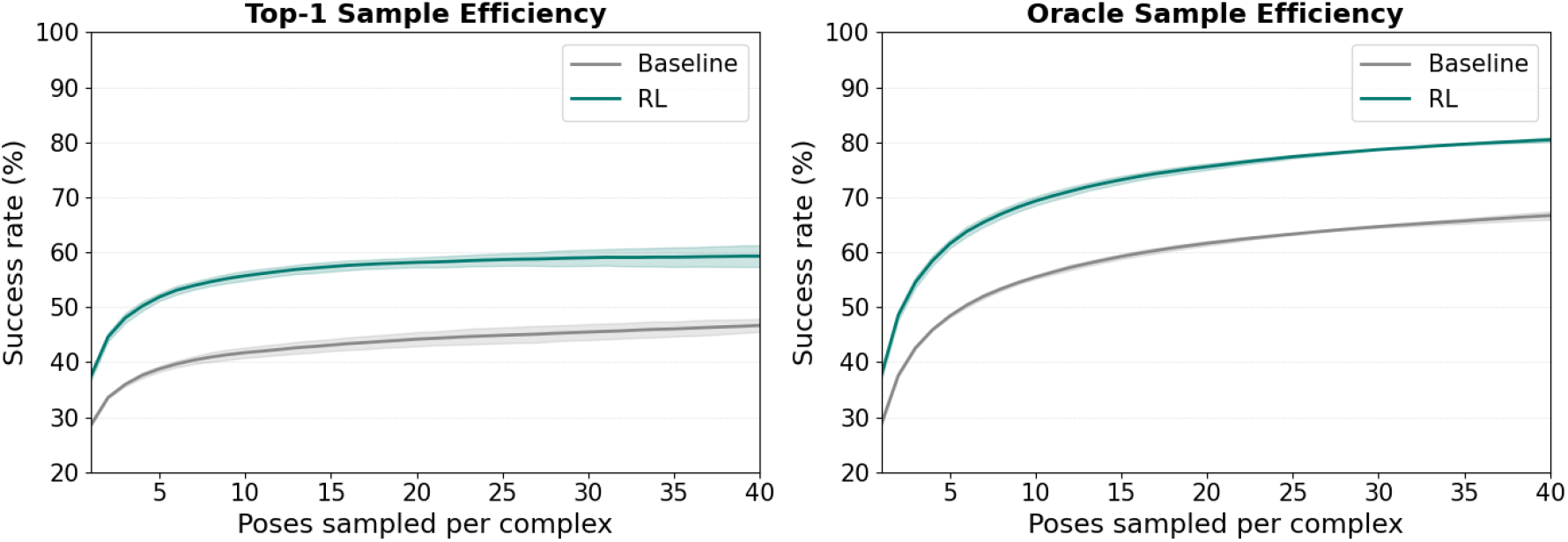
Sample efficiency curves showing success rate (RMSD ≤ 2 Å and PB-valid) as a function of poses sampled per complex. Shaded regions show standard error across runs.

## E PB-Validity Stratified by RMSD to Crystal Pose

Figure 13 shows the proportion of PB-valid poses stratified by whether the pose is within 2 Å RMSD of the crystal structure. For near-native poses (RMSD ≤ 2 Å), RL increases PB-validity from 56.6% to 79.7%, demonstrating that RL substantially improves the physical plausibility of poses that are already close to the ground truth. For poses further from the crystal structure (RMSD *>* 2 Å), PB-validity also improves from 19.9% to 40.7%. This confirms that RL does not simply trade geometric accuracy for physical validity; rather, it improves physical plausibility across both near-native and non-native poses.

**Figure 13.**
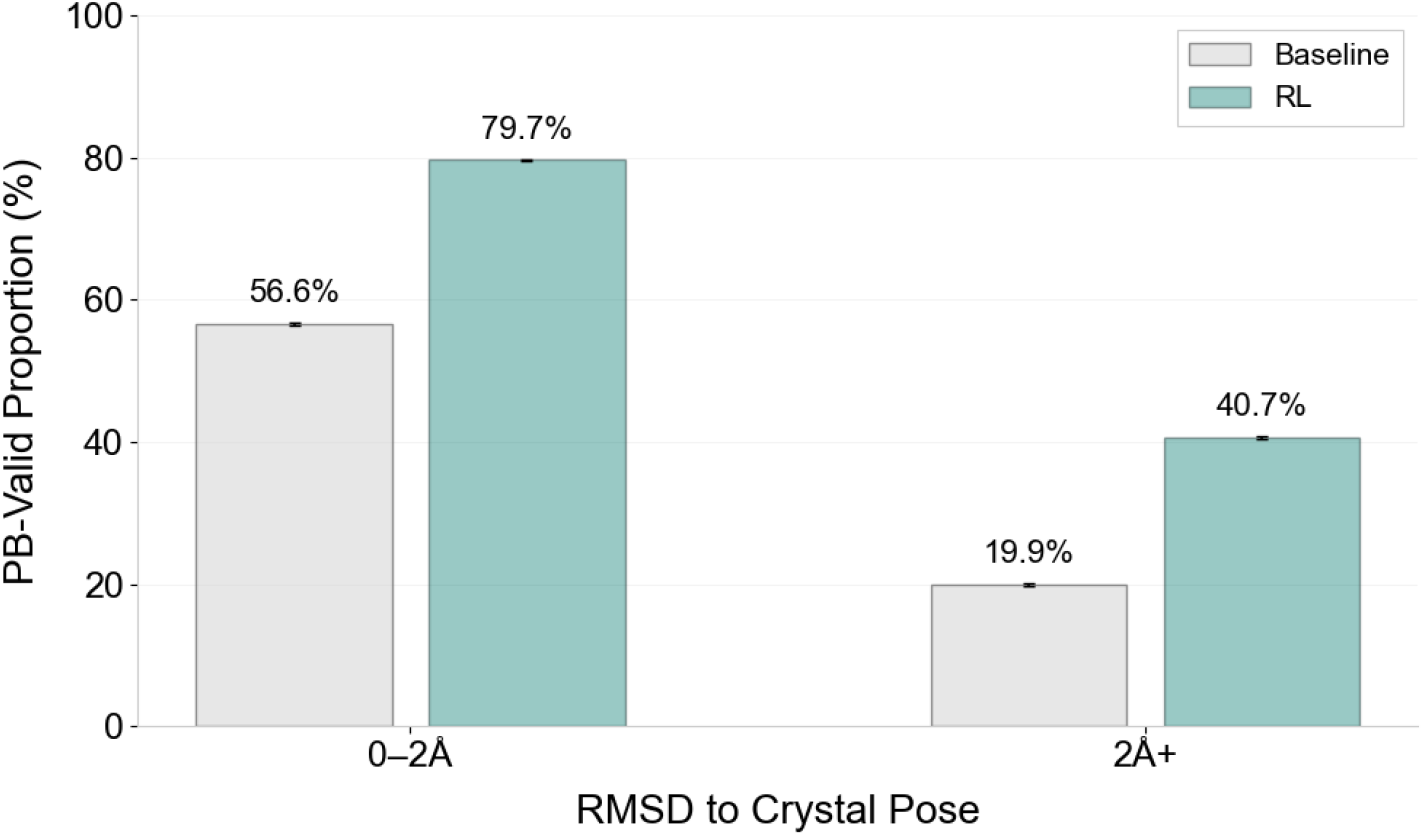
Proportion of PB-valid poses stratified by RMSD to the crystal pose. Error bars show standard error across runs.

## F Per-Complex PB-Valid Pose Comparison

Figure 14 shows the per-complex number of PB-valid poses for the overall set of poses generated for both the baseline and RL models. Each point corresponds to a single complex, with lines connecting paired baseline and RL results. Green lines indicate cases where RL generates more valid poses than the baseline, while red lines indicate cases where RL generates fewer. The violin plots show the distribution across complexes, and horizontal bars indicate the median.

**Figure 14.**
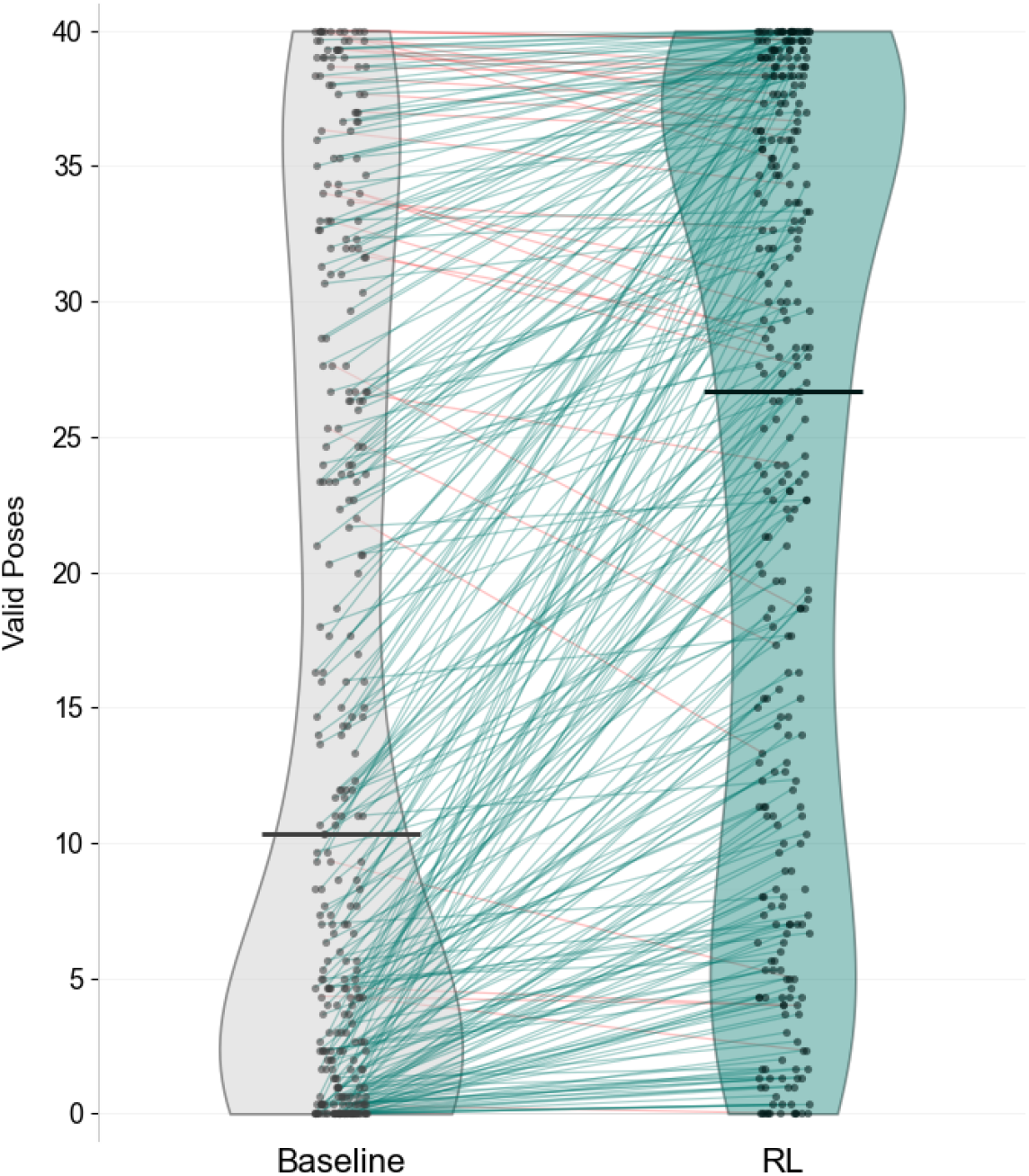
Comparison of PB-valid poses between baseline and RL. Each point represents a complex, with lines connecting paired results. Green lines indicate improvements under RL, while red lines indicate decreases. Violins show the distribution across complexes, and horizontal bars indicate medians.

## G Stratified Results by Sequence Similarity to the Training Set

To evaluate generalization performance, we stratify the PoseBusters benchmark set by sequence similarity to the training set. Results show that RL improves performance over the baseline across all similarity bins, with the strongest gains observed for the lowest similarity targets (0–30%).

**Table 5.** Success rates under increasingly strict quality criteria across sequence similarity bins (Top-1).

| Metric | 0–30% (n=109) |  | 30–95% (n=76) |  | 95–100% (n=123) |  |
| --- | --- | --- | --- | --- | --- | --- |
|  | Baseline | RL | Baseline | RL | Baseline | RL |
| RMSD $\leq 2$ Å | 56.8 | **60.5** <sup>ns</sup> | 70.1 | **72.7** <sup>ns</sup> | 73.4 | **74.3** <sup>ns</sup> |
| & PB-valid | 31.5 | **46.6** <sup>****</sup> | 50.2 | **61.5** <sup>**</sup> | 56.6 | **67.8** <sup>****</sup> |
| & $\geq 50\%$ IR | 23.8 | **37.7** <sup>****</sup> | 43.7 | **52.8** <sup>*</sup> | 49.1 | **58.3** <sup>***</sup> |
| & $\geq 75\%$ IR | 13.0 | **23.8** <sup>****</sup> | 38.1 | **39.0** <sup>ns</sup> | 36.0 | **44.4** <sup>***</sup> |

Significance assessed using McNemar’s test comparing Baseline and RL success outcomes across complexes. ns: not significant ( $p \geq 0.05$ ); \* $p < 0.05$ ; \*\* $p < 0.01$ ; \*\*\* $p < 0.001$ ; \*\*\*\* $p < 0.0001$ .

**Table 6.** Success rates under increasingly strict quality criteria across sequence similarity bins (Oracle).

| Metric | 0–30% (n=109) |  | 30–95% (n=76) |  | 95–100% (n=123) |  |
| --- | --- | --- | --- | --- | --- | --- |
|  | Baseline | RL | Baseline | RL | Baseline | RL |
| RMSD $\leq 2$ Å | 68.2 | **76.2** <sup>****</sup> | 80.5 | **87.0** <sup>**</sup> | 85.1 | **91.1** <sup>***</sup> |
| & PB-valid | 49.1 | **67.0** <sup>****</sup> | 68.0 | **82.7** <sup>****</sup> | 79.9 | **89.4** <sup>****</sup> |
| & $\geq 50\%$ IR | 43.8 | **61.7** <sup>****</sup> | 64.1 | **77.1** <sup>****</sup> | 75.1 | **84.6** <sup>****</sup> |
| & $\geq 75\%$ IR | 31.5 | **50.6** <sup>****</sup> | 56.7 | **65.4** <sup>**</sup> | 64.2 | **77.0** <sup>****</sup> |

Significance assessed using McNemar’s test comparing Baseline and RL success outcomes across complexes. ns: not significant ( $p \geq 0.05$ ); \* $p < 0.05$ ; \*\* $p < 0.01$ ; \*\*\* $p < 0.001$ ; \*\*\*\* $p < 0.0001$ .

### G.1 Minimized Structures

To evaluate the impact of minimization on generalization performance, we also stratify the PoseBusters benchmark set by sequence similarity. RL improvements are less consistent for low-similarity targets (0–30%), but remain beneficial for medium and high similarity regimes.

**Table 7.** Success rates under increasingly strict quality criteria for minimized structures (Top-1).

| Metric | 0–30% (n=108) |  | 30–95% (n=77) |  | 95–100% (n=123) |  |
| --- | --- | --- | --- | --- | --- | --- |
|  | Baseline | RL | Baseline | RL | Baseline | RL |
| RMSD $\leq 2$ Å | **70.7** <sup>ns</sup> | 69.4 | 78.4 | **84.0** <sup>*</sup> | 82.7 | **87.3** <sup>**</sup> |
| & PB-valid | **68.2** <sup>ns</sup> | 66.7 | 76.6 | **81.8** <sup>ns</sup> | 81.3 | **86.2** <sup>**</sup> |
| & $\geq 50\%$ IR | **61.4** <sup>*</sup> | 54.6 | 69.7 | **75.8** <sup>ns</sup> | 69.1 | **77.5** <sup>***</sup> |
| & $\geq 75\%$ IR | **48.1** <sup>ns</sup> | 45.4 | 53.7 | **61.5** <sup>**</sup> | 56.4 | **61.2** <sup>ns</sup> |

Significance assessed using McNemar’s test comparing Baseline and RL success outcomes across complexes. ns: not significant ( $p \geq 0.05$ ); \* $p < 0.05$ ; \*\* $p < 0.01$ ; \*\*\* $p < 0.001$ ; \*\*\*\* $p < 0.0001$ .

**Table 8.** Success rates under increasingly strict quality criteria for minimized structures (Oracle).

| Metric | 0–30% (n=108) |  | 30–95% (n=77) |  | 95–100% (n=123) |  |
| --- | --- | --- | --- | --- | --- | --- |
|  | Baseline | RL | Baseline | RL | Baseline | RL |
| RMSD $\leq 2$ Å | 76.9 | **81.5** * | 86.6 | **91.3** * | 88.1 | **93.0** ** |
| & PB-valid | 75.3 | **79.6** * | 85.3 | **90.5** * | 88.1 | **93.0** ** |
| & $\geq 50\%$ IR | 73.1 | **73.5** <sup>ns</sup> | 81.4 | **86.1** <sup>ns</sup> | 82.4 | **88.1** ** |
| & $\geq 75\%$ IR | 63.6 | **67.3** <sup>ns</sup> | 73.6 | **73.6** <sup>ns</sup> | 71.3 | **79.7** *** |

Significance assessed using McNemar’s test comparing Baseline and RL success outcomes across complexes. ns: not significant ( $p \geq 0.05$ ); \* $p < 0.05$ ; \*\* $p < 0.01$ ; \*\*\* $p < 0.001$ ; \*\*\*\* $p < 0.0001$ .

## H Overlap in Success Sets Across Sequence Identity Groups

Figures 15 to 17 show how the success sets of the original DiffDock-Pocket and the RL-trained DiffDock-Pocket model overlap across three sequence identity groups. As the success criteria becomes more strict, the success sets shrink and the overlap typically decreases. This creates greater divergence between the models, meaning one model succeeds on complexes where the other fails. Across all groups, the divergence increasingly favours RL under strict criteria. This effect is strongest for low sequence identity targets, which are the most out of distribution.

**Figure 15.**
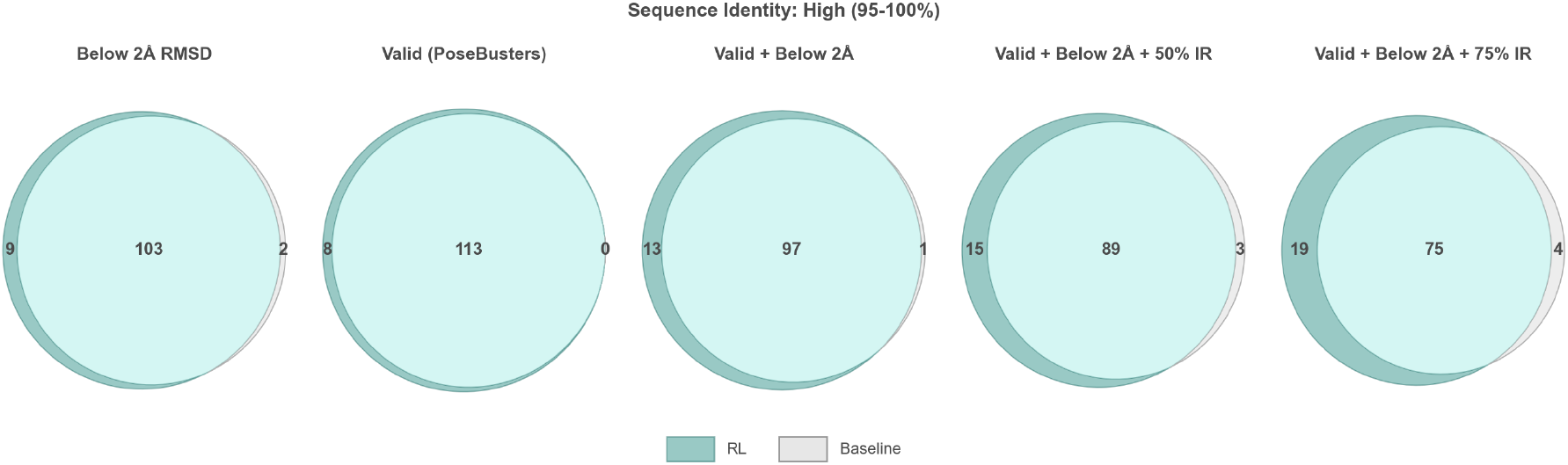
Overlap in success sets between the baseline DiffDock-Pocket model and the RL trained model for high sequence identity targets in the range 95%–100%. *N* = 123.

**Figure 16.**
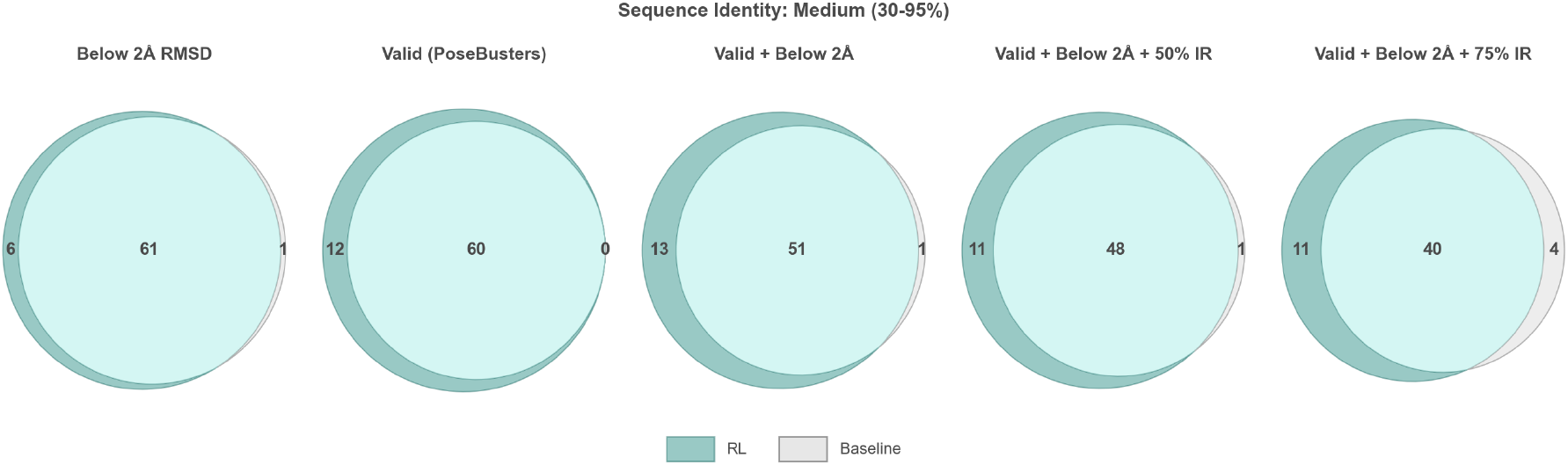
Overlap in success sets between the baseline DiffDock-Pocket model and the RL trained model for medium sequence identity targets in the range 30%–95%. *N* = 76.

For sequence identity in the range 95%–100%, many successes are shared between the two models across all criteria. This pattern is consistent with an in-distribution regime where the baseline already performs well. As the success criteria become more strict, the sets shrink but most successes remain shared. RL still contributes additional successes, and these appear most clearly when success requires PB-validity and RMSD ≤ 2Å as well as recovery of 75% of ground-truth interactions.

**Figure 17.**
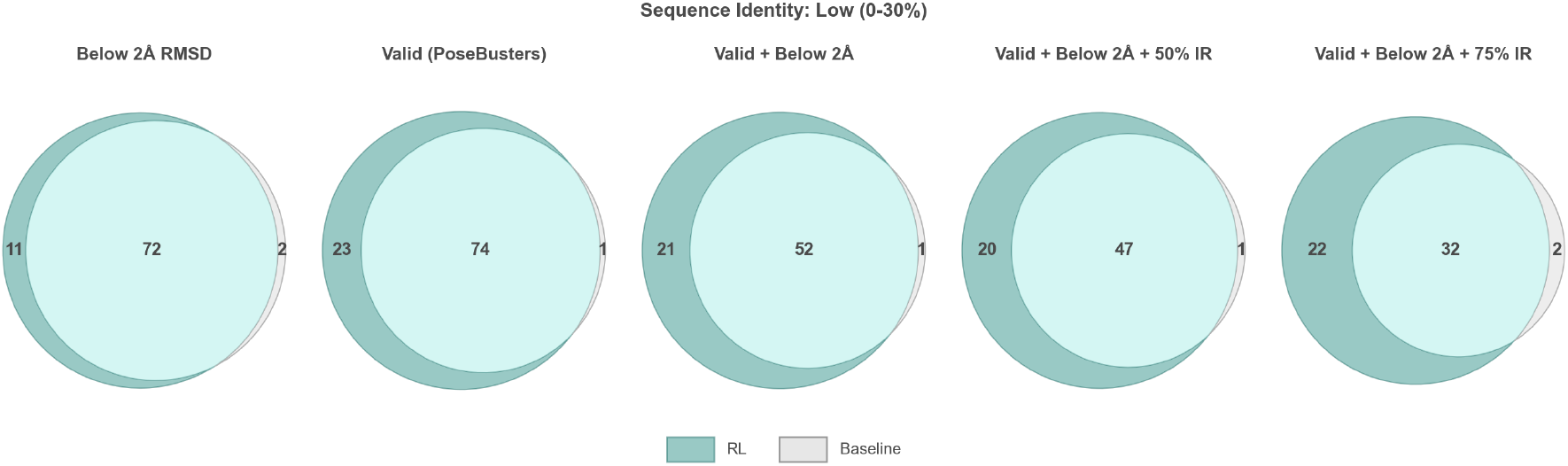
Overlap in success sets between the baseline DiffDock-Pocket model and the RL trained model for low sequence identity targets in the range 0%–30%. *N* = 109.

For sequence identity in the range 30%–95%, when success is defined by PB validity and RMSD below 2 Å, the overlap decreases and model specific successes become more frequent. This indicates that the harder requirement is not simply generating a near native pose or generating a plausible pose in isolation, but generating a pose that satisfies both at the same time. RL improves this joint success more consistently than the baseline, and the gap becomes more visible once interaction recovery thresholds are included.

For sequence identity in the range 0%–30%, the success sets diverge substantially as the criteria become more strict. The baseline retains relatively few successes once PB validity and RMSD below 2 Åare required simultaneously, and it contracts further when interaction recovery thresholds are added. In contrast, RL preserves a much larger successful set under these strict criteria, producing many RL only successes. This widening divergence supports the conclusion that reinforcement learning most strongly improves performance on the most out of distribution targets, particularly when success requires joint near nativeness, physical plausibility, and interaction recovery.

## I Training hyperparameters

**Table 9.**
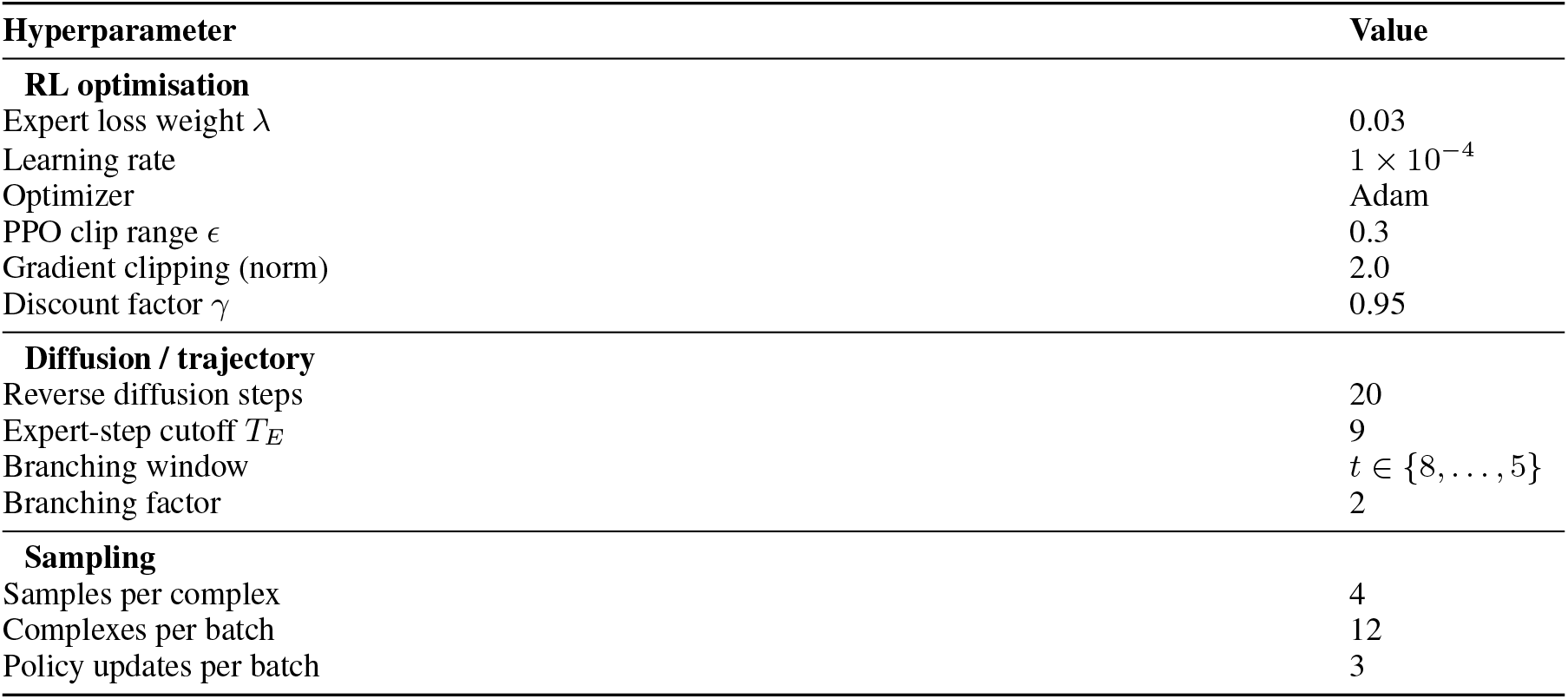
Training hyperparameters for reinforcement learning fine-tuning of DiffDock-Pocket.

